# Ruler elements in chromatin remodelers set nucleosome array spacing and phasing

**DOI:** 10.1101/2020.02.28.969618

**Authors:** Elisa Oberbeckmann, Vanessa Niebauer, Shinya Watanabe, Lucas Farnung, Manuela Moldt, Andrea Schmid, Patrick Cramer, Craig L. Peterson, Sebastian Eustermann, Karl-Peter Hopfner, Philipp Korber

## Abstract

Arrays of regularly spaced nucleosomes dominate chromatin and are often phased by alignment to reference sites like active promoters. How the distances between nucleosomes (spacing), and between phasing sites and nucleosomes are determined remains unclear, and specifically, how ATP dependent chromatin remodelers impact these features. Here, we used genome-wide reconstitution to probe how *Saccharomyces cerevisiae* ATP dependent remodelers generate phased arrays of regularly spaced nucleosomes. We find that remodelers bear a functional element named the ‘ruler’ that determines spacing and phasing in a remodeler-specific way. We use structure-based mutagenesis to identify and tune the ruler element residing in the Nhp10 and Arp8 modules of the INO80 remodeler complex. Generally, we propose that a remodeler ruler regulates nucleosome sliding direction bias in response to (epi)genetic information. This finally conceptualizes how remodeler-mediated nucleosome dynamics determine stable steady-state nucleosome positioning relative to other nucleosomes, DNA bound factors, DNA ends and DNA sequence elements.

---

Nuclear DNA is packaged into chromatin based on a repeating building block, the nucleosome core particle (NCP; (Kornberg, 1974; Olins and Olins, 1974)), where 147 base pairs (bp) of DNA are wound around a histone protein octamer (Kornberg and Lorch, 1999; Luger et al., 1997; Olins and Olins, 2003). Packaging by nucleosomes orchestrates all genomic processes (Lai and Pugh, 2017).

Nucleosomes mainly occur in regular arrays where they are aligned to each other such that the lengths of linker DNA between NCPs are about constant within an array. Linker lengths may vary among arrays in the same cell (Baldi et al., 2018b; Chereji et al., 2018; Ocampo et al., 2016; Valouev et al., 2011) and differ on average between cell types and species (van Holde, 1989). Arrays are often phased, i.e., aligned relative to a genomic reference point. A combination of both *in vivo* studies (Ganapathi et al., 2011; Hartley and Madhani, 2009; Kubik et al., 2018; Tsankov et al., 2011; van Bakel et al., 2013; Yan et al., 2018; Yarragudi et al., 2004) and *in vitro* reconstitutions (Krietenstein et al., 2016) indicated that these genomic alignment points or “barriers” often reflect the binding of abundant, sequence-specific DNA binding proteins, like the general regulatory factor (GRFs) Reb1, Abf1, or Rap1 in budding yeast or other architectural factors like CTCF in mammals (Wiechens et al., 2016) or Phaser in flies (Baldi et al., 2018a).

Throughout eukaryotes, phased arrays are prominent at active promoters. Nucleosome-depleted regions (NDRs) at the core promoter are flanked by arrays that begin with the so called +1 nucleosome close to the transcription start site (TSS) and cover the gene body (Baldi et al., 2020; Lai and Pugh, 2017). This organization is important for transcription fidelity as mutants with impaired array phasing show aberrant transcription initiation (Challal et al., 2018; Hennig et al., 2012; Kubik et al., 2019; Pointner et al., 2012; Smolle et al., 2012). While nucleosome arrays are likely the most pervasive and longest known chromatin organization, their generation is still not explained. Specifically, regular spacing requires fixed distances between nucleosomes, and phasing requires a fixed distance between array and reference point. What sets these distances?

*In vivo* and *in vitro* data suggest that ATP dependent chromatin remodeling enzymes (remodelers) are key to the answer. Remodelers are conserved in eukaryotes (Flaus et al., 2006) and mobilize, reconfigure, or disassemble/reassemble nucleosomes upon ATP hydrolysis (Clapier and Cairns, 2009; Clapier et al., 2017). They are subdivided into the SWI/SNF, ISWI, CHD, and INO80 families, according to their main ATPase sequence features. Besides the core ATPase, remodelers often contain additional domains and subunits that bind the nucleosome, regulate activity and targeting, and convert their DNA tracking activity into the remodeler-specific chemo-mechanical reaction. For example, nucleosome disassembly is accomplished only by SWI/SNF family members and histone exchange only by INO80 family members, while nucleosome sliding is catalyzed by most remodelers. Particularly relevant for array generation is an ATP-dependent nucleosome spacing activity, by which some remodelers convert irregular arrays into arrays of regularly spaced nucleosomes. Remodelers of the ISWI, CHD, and INO80 (Ito et al., 1997; Tsukiyama et al., 1999; Udugama et al., 2011; Varga-Weisz et al., 1997), but not of the SWI/SNF family, show spacing activity. This activity was suggested to rely on a length-sensor mechanism (Yang et al., 2006; Zhou et al., 2018) where nucleosome sliding rate is regulated by linker DNA length. Sliding one nucleosome back and forth between two other nucleosomes, with a linker length-dependent velocity, would center a nucleosome at steady state when both flanking linkers have the same length.

While the length-sensor mechanism may equalize linker lengths and thereby generate spacing distance *regularity*, it does not by itself determine spacing distance *length* in absolute terms. This would reciprocally depend on nucleosome density. However, spacing *in vivo* (Gossett and Lieb, 2012; Hennig et al., 2012; van Bakel et al., 2013), as well as generated *in vitro* (Lieleg et al., 2015; Zhang et al., 2011), remained constant despite changes in nucleosome density. This was called “active packing” (Zhang et al., 2011) or “clamping” (Lieleg et al., 2015), but it remained unclear if remodeler or nucleosome features led to such density-independent spacing. Structural studies suggested that the yeast ISW1a remodeler contacts a neighboring nucleosome and may set the linker length by a “protein ruler” (Yamada et al., 2011). Two ISWI family remodelers, yeast ISW1a and ISW2, each generated regular arrays aligned at DNA-bound Reb1 or Abf1 *in vitro*, but with different spacing at the same nucleosome density (Krietenstein et al., 2016). This points towards a remodeler-specific linker length determining ruler mechanism. Also suggestive of a built-in ruler, INO80 required a minimum linker length for nucleosome sliding (Zhou et al., 2018) and recognized linker DNA via a structural module that was important for sliding (Knoll et al., 2018).

The ruler metaphor may indeed describe a remodeler mechanism that measures and sets the phasing and spacing distances of arrays. However, so far it is mainly suggestive and has to be substantiated in molecular terms. This would be exceedingly convoluted *in vivo* but requires a defined system that allows to assay the generation of phased regular arrays by remodelers and to dissect if and how a ruler mechanism is at work. Are there rulers within some or all remodelers with spacing activity? Are linker length vs. distance to barrier determined in the same or different way? Are rulers autonomous or does the outcome depend on nucleosome density or underlying DNA sequence? Ultimately, is it possible to tune a ruler, i.e., can a remodeler be mutated to generate arrays with altered spacing and/or phasing distances?

Here, we used genome-wide *in vitro* chromatin reconstitution with purified remodelers ((Krietenstein et al., 2016), accompanying paper Oberbeckmann & Krietenstein et al.) to answer these questions. All yeast remodelers with spacing activity, ISW1a, ISW2, Chd1, and INO80 have rulers that are largely autonomous regarding underlying DNA sequence but some may respond to nucleosome density. Remodeler-specific rulers mechanistically explain earlier *in vivo* observations. Structure-guided mutations in recombinant INO80 complexes led to shorter or longer spacing and phasing distances and showed that these quantities may be uncoupled. Finally, we propose a model how remodeler rulers position nucleosomes by regulating sliding direction bias according to (epi)genetic information in the nucleosome environment.

## Results

### Defined genome-wide chromatin reconstitution system with varying nucleosome densities

To assess array generation by remodelers in a biochemically defined way, we used our genome-wide chromatin reconstitution system with purified components (Figure 1A, (Krietenstein et al., 2016)) including recombinant INO80 complex (accompanying paper Oberbeckmann & Krietenstein et al.) and recombinant Chd1 (Farnung et al., 2017). Briefly, genomic plasmid libraries were reconstituted with *Drosophila* embryo histone octamers into nucleosomes by salt gradient dialysis (SGD). SGD chromatin was incubated with ATP, purified yeast remodelers (Figure S1A), and the barrier Reb1 or the restriction enzyme BamHI, which generates double strand breaks (DSBs) that also amount to nucleosome positioning barriers (accompanying paper Oberbeckmann & Krietenstein et al.). Resulting nucleosome patterns were analyzed by MNase-seq. The effective histone-to-DNA mass ratio during SGD was varied from 0.2 to 0.8 yielding low, medium and high nucleosome densities reflected in increasingly extensive MNase-ladders at the same MNase digestion conditions (Figure 1B). Nucleosome density variation was instrumental to distinguish if linker lengths and phasing distances depended on nucleosome density and/or remodeler features.

**Figure 1.**
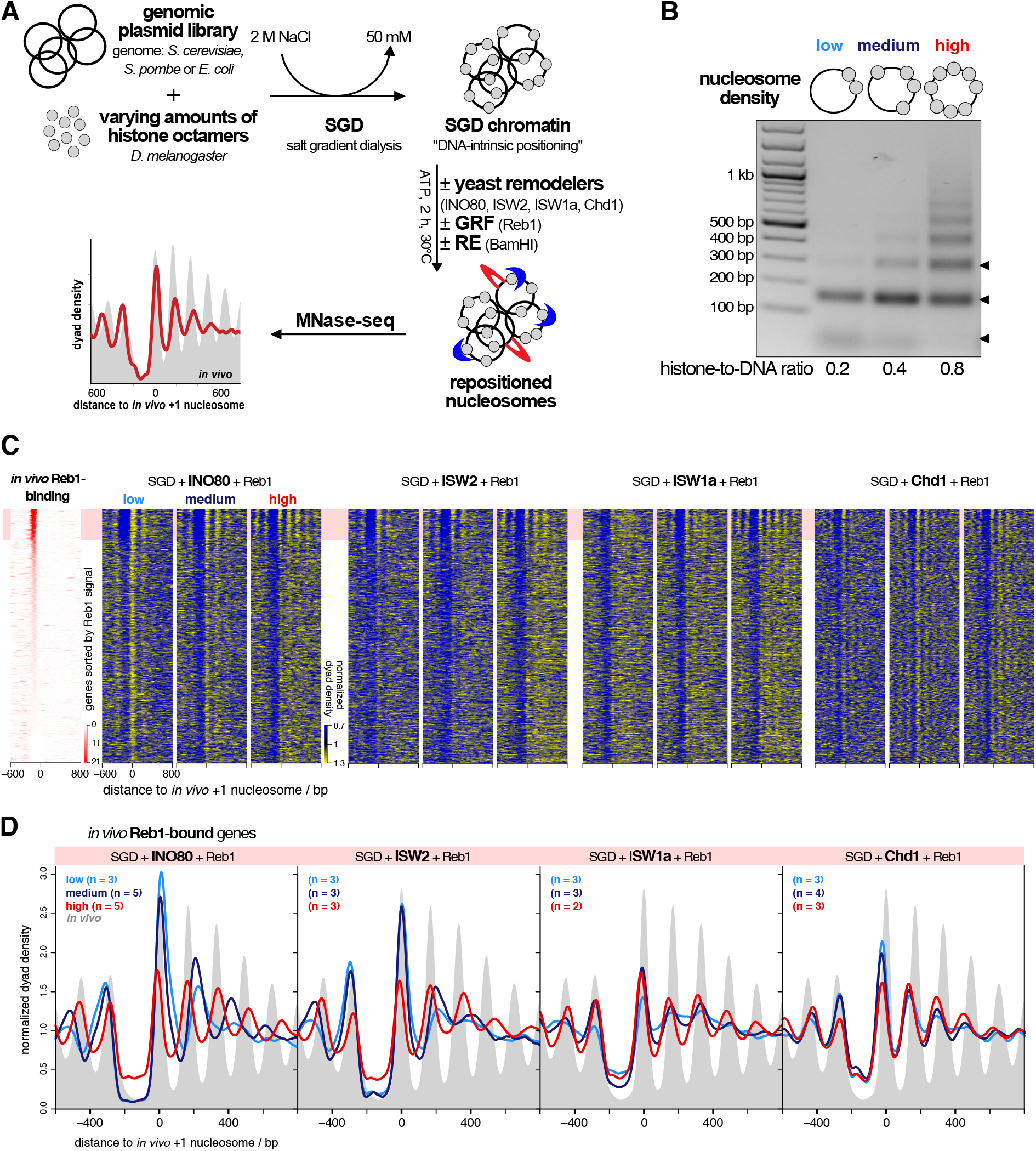
Rebl-guided nucleosome positioning *in vitro* by individual remodelers at varying nucleosome density. **(A)** Overview of genome-wide *in vitro* reconstitution system. **(B)** Comparison of SGD chromatin reconstituted at indicated histone-to-DNA ratios. DNA fragments after MNase-digest under the same conditions were resolved by agarose gel electrophoresis. Arrow heads on right point to subnucleosomal, mono- and dinucleosomal fragments (bottom to top). **(C)** Heat maps of MNase-seq data for SGD chromatin of the indicated nucleosome density and after incubation with indicated remodelers and Reb1. Chd1 refers to the Chd1/FACT complex. Heat maps are aligned at *in vivo* +1 nucleosome positions and sorted according to decreasing (top to bottom) anti-Reb1 SLIM-ChIP score (Gutin et al., 2018) shown in leftmost heat map. Horizontal red shading highlights genes with strong *in vivo* Reb1-binding in their promoters. Merged data of replicates are shown, individual replicates in Figure S1B,C. **(D)** Composite plots for MNase-seq data averaged over the indicated number of replicates (n) as in panel B but only for genes highlighted in red in panel B.

### INO80, ISW2, ISW1a and Chd1, but not Fun30 align regular arrays at the barrier Reb1

We tested all yeast remodelers with known spacing activity, INO80, ISW2, ISW1a and Chd1 (Krietenstein et al., 2016; Lusser et al., 2005; Stockdale et al., 2006; Torigoe et al., 2013; Tsukiyama et al., 1999; Udugama et al., 2011) as well as the Fun30 remodeler, for which it was unclear if it has spacing activity (Awad et al., 2010). INO80, ISW2, ISW1a and Chd1, each in combination with Reb1, generated phased regular arrays at promoters with Reb1 sites (red shaded top of heat maps in Figure 1C), while Fun30 did not (Figure S1B). This clarifies that Fun30 does not have regular array generation and alignment activity. Previously, Chd1 purified from budding yeast did not show much effect in genome-wide reconstitutions (Krietenstein et al., 2016). This was maybe due to full-length Chd1 tending to aggregate *in vitro*, which is why truncated Chd1 constructs were often used (McKnight et al., 2011; Patel et al., 2011). Here, we leveraged our finding that recombinant full-length Chd1 is stabilized in complex with recombinant FACT complex (Farnung et al., 2017) and achieved *in vitro* array generation and alignment also by Chd1.

The heat map patterns (Figure 1C) and even more the corresponding composite plots for the Reb1-bound genes only (Figure 1D) suggested that the distance of arrays to the barrier Reb1 as well as the linker lengths varied with nucleosome density in a remodeler-specific way. For all remodelers with spacing activity, array extent increased with growing density, consistent with greater nucleosome availability and processive spacing activity. Array extent at high density was larger than in our previous reconstitutions (Krietenstein et al., 2016), i.e., we achieved higher densities here. Adding more remodeler after half of the incubation time did not change the array distances of resulting patterns confirming non-limiting remodeling activity and steady state conditions (Figure S1C).

### Remodelers set phasing and spacing distances symmetrically around barriers

To better assess distances to barrier (phasing) and linker lengths (spacing), we aligned the MNase-seq data for each remodeler/barrier/density combination to either *in vivo* Reb1 sites or BamHI sites (Figure 2A). For each replicate (Figure S2A-C), we called nucleosome peaks and determined the distances to barrier and linker lengths as defined in Figure 2B.

**Figure 2.**
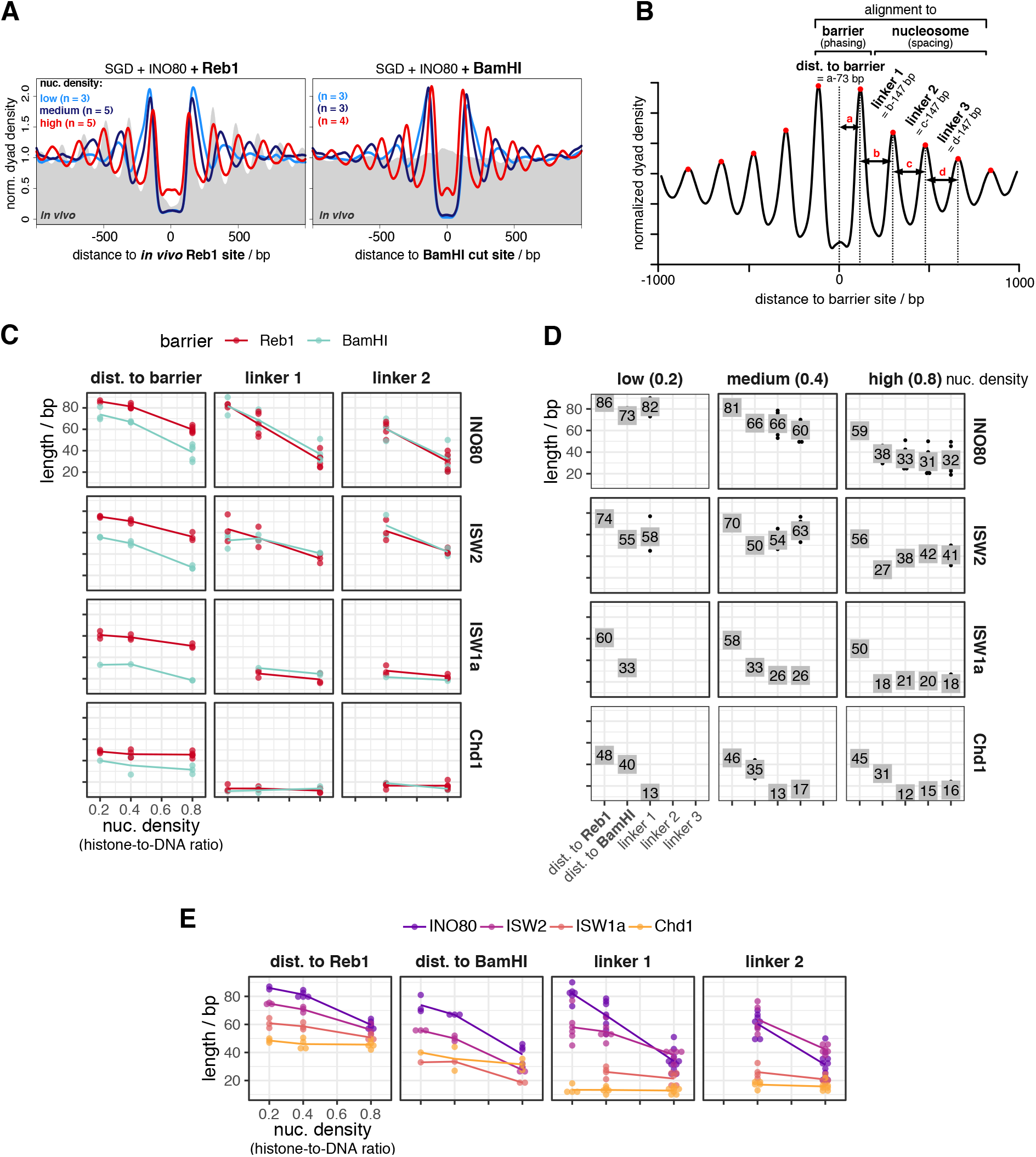
Quantification of barrier-aligned nucleosome array features depending on barrier, remodeler and nucleosome density. **(A)** Composite plots of same MNase-seq data for INO80 as in Figure 1D but aligned at anti-Reb1 SLIM-ChlP-defined Reb1 sites (left), or at BamHI sites (right) of SGD chromatin reconstituted at the indicated nucleosome densities and incubated with INO80 and BamHI. **(B)** Scheme defining array features quantified from barrier-aligned composite plots as in panel A. **(C) – (D)** Array feature values for the indicated combinations of barrier, remodeler and nucleosome density plotted in different ways allowing comparison between barriers (especially panel C), values (especially panel D) and remodelers (especially panel E). Chd1 refers to the Chd1/FACT complex. Panel D and Figure S2A-C show individual replicates, panels C and E replicate averages.

All remodelers symmetrically aligned regular arrays to BamHI sites, which are palindromic and therefore inherently symmetrical, and most of them also to Reb1 sites (Figures 2A, S2A,B) regardless of site orientation and position relative to genes (groups 1 to 3; Figure S3A,B). However, if INO80 aligned arrays at promoter Reb1 sites (groups 1 to 3, Figure S3A, accompanying paper by Oberbeckmann & Krietenstein et al.), nucleosome occupancy (peak height) was higher over genic versus non-genic regions at low and medium nucleosome density leading to asymmetric patterns with regard to peak heights in groups 1 and 2. Reb1 site orientation had no effect (group 1 vs. 2). This asymmetry in nucleosome occupancies reflected that positioning of +1 nucleosomes, per definition the first nucleosomes downstream of transcription start sites, i.e. at gene starts, was not only guided by Reb1 bound to promoter sites but also synergistically by underlying DNA shape features (accompanying paper Oberbeckmann & Krietenstein et al.). We recapitulated here that INO80 was able to position *in vivo*-like +1 nucleosomes in the absence of a barrier at low and medium densities (Figure S2C,D). This synergism between Reb1- and DNA shape-guided +1 positioning at low and medium density resulted in higher occupancy at the +1 nucleosomes, which are alignment points for +2 nucleosomes and so on. Therefore, all array peaks over genes were higher than their counterparts over non-genic regions. However, such synergism was not seen at high density where *in vivo*-like +1 nucleosomes positioning by INO80 alone was much less pronounced (Figure S3C,D). This inability was not due to a general inability of INO80 to slide densely packed nucleosomes as INO80 could generate Reb1-aligned arrays at these high nucleosome densities, too (Figures 1C,D, 2A, S2A,B). Nonetheless, this activity was apparently incompatible with or dominant over DNA shape-guided nucleosome positioning (see Discussion). This showed again that our here generated high nucleosome density was higher than the nucleosome density used previously (Krietenstein et al., 2016), otherwise *in vivo*-like +1 nucleosome positioning by INO80 would not have been clearly observed in our earlier study.

In this context, we also tested if Fun30 positions *in* vivo-like +1/-1 nucleosomes on its own, but it did not (Figure S3D).

In contrast to nucleosome peak *heights*, nucleosome peak *positions* and therefore corresponding phasing and spacing distances were not significantly affected across groups 1 to 3 for all remodelers, including INO80 (Figure S3B). Therefore, all remodelers symmetrically generated phasing and spacing distances at Reb1 and BamHI sites, which warranted averaging over the up- and downstream values. Resulting values were plotted in different ways to facilitate multi-dimensional comparisons (Figure 2C-E). As all remodelers generated linker lengths independently of the barrier type, we combined linker length values for both barriers (Reb1 and BamHI, Figure 2C). Linker length determination relied on nucleosome peak calling, which was often not possible beyond the −1/+1 nucleosomes at low nucleosome density (Figure S2A), so that linker length data for low density conditions were more sparse, even absent for ISW1a.

### Remodeler-specific rulers set spacing in a density-independent or –dependent way

To compare spacing generated by different remodelers at different nucleosome densities, we focused on the averaged length of linker 1 (Figure 2B), which was most accessible across all nucleosome densities. Chd1 generated the shortest (12-13 bp) and ISW1a a bit longer (21-26 bp) linker 1 lengths without significant effects by nucleosome density (Figure 2D,E). ISW2 generated rather constant spacing (54-58 bp) at low and medium but tighter spacing (38 bp) at high density. For INO80, linker lengths steadily increased with decreasing density from 33 to 82 bp. We concluded that linker lengths and their dependencies on nucleosome density were remodeler-specific and interpreted this as follows. Spacing activity of a remodeler has two aspects. On the one hand, the remodeler equalizes linker lengths leading to regularity in arrays, which is the classical definition of spacing activity (Ito et al., 1997; Varga-Weisz et al., 1997). On the other hand, the resulting linkers have a certain length. In our purified system, this may either be determined by nucleosome density and/or by a remodeler-intrinsic feature. Following (Yamada et al., 2011), we call a remodeler feature that sets nucleosome spacing a “ruler”. We use this term also for the feature that sets the distance to barriers (see below). Indicative for a remodeler ruler is remodeler-specific clamping, i.e., if constant spacing is generated at different nucleosome densities (= clamping) and different remodelers generate different spacing (= remodeler-specific), which shows that spacing depends on remodeler-intrinsic and not nucleosome-intrinsic properties (Lieleg et al., 2015). We saw remodeler-specific clamping for Chd1 at all, for ISW1a at high versus medium and for ISW2 at medium versus low densities (Figure 2C-E). As none of the remodelers with spacing activity can disassemble nucleosomes (Clapier and Cairns, 2009) and thereby affect nucleosome density, their rulers can only set their respective linker lengths if these are shorter than or equal to the density-determined linker length at equidistant nucleosome distribution. Accordingly, Chd1 and ISW1a set their ruler-specified linker lengths at all and ISW2 at medium and low densities. ISW2 had to generate shorter linkers at high density and INO80 either did not have a ruler or the ruler responded to changes in nucleosome density.

*In vitro* mononucleosome assays suggested that INO80 requires at least 40 bp of nucleosome-free DNA for nucleosome sliding (Zhou et al., 2018), while it generated 30 bp linkers in tri-nucleosomes (Udugama et al., 2011). Here, at high nucleosome density, INO80 generated linkers of about 33 bp consistent with previous observations. We tried to enforce even tighter spacing by increasing nucleosome density. This did not decrease spacing and phasing distances but peak heights (Figure S2B,C), probably due to increased aggregation without effective increase in nucleosome density of soluble chromatin.

### Remodeler type, barrier type and nucleosome density determine distance to barrier

The findings for the distance to barrier were more complex than for lengths of linker 1 (Figure 2C-E). First, the distance to barrier depended on the barrier type (Figure 2C). It was always longer for Reb1 than for BamHI generated DNA ends, with the largest difference for ISW1a and the smallest for Chd1. The DNA footprint size of *S. cerevisiae* Reb1 is not known, possibly 20 bp as for the *S. pombe* Reb1 DNA binding domain (Jaiswal et al., 2016). This would contribute 10 bp to the distance to barrier (Figure 2B) and could explain the differences between distance to Reb1 vs. BamHI sites for Chd1, but not for the other remodelers. Therefore, INO80, ISW2 and ISW1a, but not Chd1, aligned nucleosomes differently at Reb1 versus at DSBs.

Second, the distance to DNA ends was mostly similar to linker lengths for INO80, ISW2 and ISW1a, arguing that these remodelers, but not Chd1, used a DNA end in a similar way as a neighboring nucleosome for nucleosome alignment.

Third, distances to barriers depended on nucleosome density in a similar way as linker lengths for all remodelers but INO80, where distances to both barriers varied less between low and medium density than linker length.

We concluded that there are remodeler-specific differences in how a nucleosome is positioned next to another nucleosome versus next to a barrier like Reb1 versus next to a DNA end and how this depends on nucleosome density. This is again a clear case of different remodelers generating different nucleosome positioning, although starting from the same SGD chromatin, which argues for remodeler-specific rulers governing nucleosome positioning.

### Remodelers differ in processivity of nucleosome positioning

All remodelers generated similar lengths of linker 1 to linker 3 at high density (Figure 2D), which we interpreted as processive spacing activity along the arrays as long as nucleosomes were sufficiently provided. At low density, ISW2, Chd1 and especially INO80 still generated high +1/-1 nucleosome peaks (Figure S2A), in contrast to ISW1a, for which these peaks were less pronounced and +2/2 nucleosome peaks could not be discerned. We suggest that ISW1a is less processive than other remodelers in bringing nucleosomes next to barriers at low densities.

### Remodelers generate similar arrays on all but more effectively on eukaryotic DNA sequences

The same linker lengths in arrays at BamHI and Reb1 sites (Figure 2C), at Reb1 sites in groups 1 to 3 and the symmetry of nucleosome distances to Reb1 sites in groups 1 to 3 (Figure S3A,B) suggested that remodeler rulers position nucleosomes independently of DNA sequence flanking the barriers. Nonetheless, there are evolved DNA features at promoters, especially for INO80 (accompanying paper Oberbeckmann & Krietenstein et al.), that affected occupancies (peak heights, not positions, Figure S3A), which may also be true for evolved nucleosome-favoring dinucleotide periodicities (Satchwell et al., 1986) in gene bodies.

To rigorously disentangle these contributions, we tested the remodeler/barrier/density combinations also with SGD chromatin of *S. pombe* and *E. coli* genomic plasmid libraries (Figures 1A, 3A,B, S4A), including the steady state control (Figure S4B). We did not observe substantial differences in spacing/phasing distances on these genomes for all remodelers, but some replicates, especially at medium and low density, showed lower relative occupancies for the *E. coli* genome.

**Figure 3.**
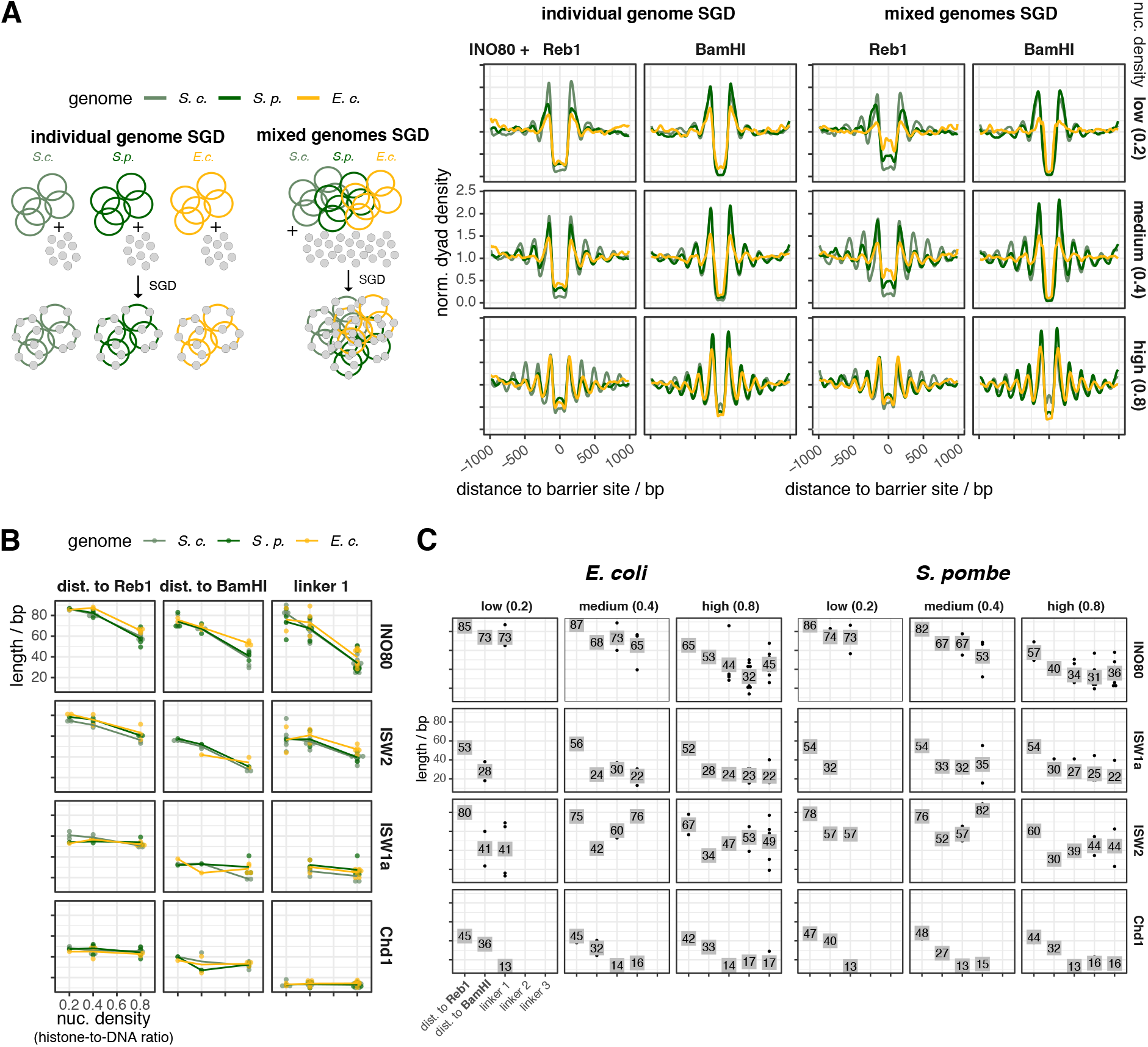
Yeast remodelers generate arrays on heterologous genomes with same spacing and phasing distances as on the yeast genome. **(A)** Left: Schematic showing SGD reconstitution with individual or mixed genomes. Right: Composite plots as Figure 2A but for the indicated barriers and either individual (left) or mixed (right) genomes. Reb1 sites were called by PWM. **(B)** and **(C)** As Figures 2C,D, respectively, but for the indicated genomes. Individual replicates in Figure S4.

We concluded that all remodelers align arrays at Reb1 or DSBs regardless of the underlying sequence. Nonetheless, they are more effective in terms of relative occupancies on eukaryotic genomes, likely due to dinucleotide periodicities (Zhang et al., 2009).

### INO80 complexes mutated in the Arp8 and/or Nhp10 module

It was unexpected that the clamping criterion did not clearly show a ruler for INO80 (Figure 2C-E), because the INO80 structure suggested modules that bind extranucleosomal DNA and could serve as ruler (Knoll et al., 2018). To clarify, we took advantage of the biochemical accessibility of our recombinant INO80 preparation, the modular INO80 composition and the high-resolution structures (Eustermann et al., 2018; Knoll et al., 2018) to generate candidate mutations that may tune and thereby reveal INO80’s ruler.

The INO80 complex has two modules with a likely role in ruler function. First, the Arp8 module consisting of N-Actin, Arp8, Arp4, Taf14 and Ies4 (Figure 4A). It binds the Ino80 main ATPase HSA domain, which is structured as a long helix with a kink that subdivides it into the HSAα1 and HSAα2 part (Knoll et al., 2018). Both bind to extranucleosomal DNA, and mutating DNA contacting lysine residues in HSAα1 or HSAα2 to glutamines (HQ1 and HQ2 mutant, respectively, Figure 4B,C,D) impaired, and combining both mutations (HQ1/2 mutant) abolished mononucleosome centering activity (Knoll et al., 2018).

**Figure 4.**
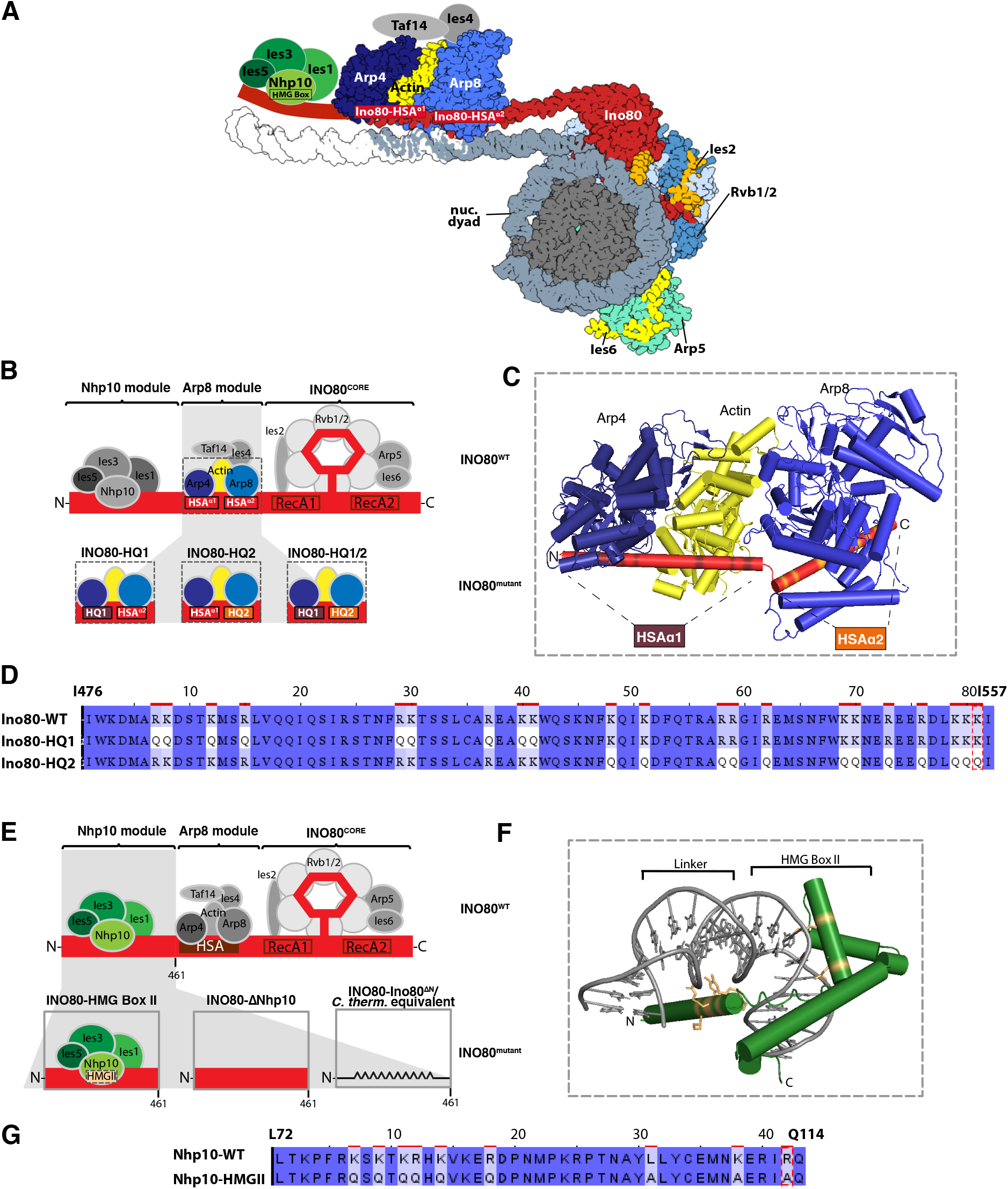
Construction of INO80 mutant complexes. **(A)** Structure-based (Eustermann et al., 2018; Knoll et al., 2018) model of a nucleosome bound by the INO80 complex with indicated subunits. Nhp10 module, Taf14 and Ies4 organization is assumed. **(B)** Schematic of INO80 complex submodule and subunit organization (top). Zoom into Arp8 module showing three mutant versions (bottom). **(C)** Cylindrical representation of the Arp8 module structure showing mutated residues of Ino80 HSA domain (highlighted in brown and orange). **(D)** Sequence alignment showing mutated residues in Ino80-HQ1 and –HQ2 mutants. **(E)** Schematic of INO80 complex organization as in panel B (top) but zoom into Nhp10 module (bottom) showing three mutant versions. **(F)** Model of Nhp10 HMG box-like and Linker region (residues 62-172) based on TFAM structure (pdb 3tq6). **(G)** Sequence alignment showing mutated residues in Nhp10-HMGII mutant.

The second, Nhp10 module, binds the Ino80 ATPase N-terminus, and contains the HMG box Nhp10 subunit, along with Ies1, Ies3 and Ies5 (Figure 4A,E). This module is species-specific and affects the processivity and extranucleosomal DNA requirements in mononucleosome sliding assays (Zhou et al., 2018).

Calculating a homology model for Nhp10 based on another HMG box protein, TFAM (Ngo et al., 2014), we inferred and mutated amino acid residues putatively involved in Nhp10-DNA interactions (HMGII mutant, Figure 4F,G). These mutations were also combined with the HQ1 or HQ2 mutants (HMGII-HQ1 and HMGII-HQ2). Further, we prepared recombinant INO80 complex without any Nhp10 module subunits (ΔNhp10 mutant, no truncation of the Ino80 ATPase N-terminus) or a version where the Ino80 ATPase lacked residues 1-461 (INO80^ΔN^ mutant), which removes the assembly platform for the Nhp10 module (Figure 4A).

### INO80 mutant complexes reveal a multilayered ruler

All mutant complexes were assayed like the wild type (WT) INO80 complex (Figures 5A-E, 6A-D, S1A, S5A,B). WT INO80 was assayed again alongside with matching SGD chromatin. Comparing these replicates (Figure 5C) with previous values for WT INO80 (Figure 2D) reflected variability in preparing SGD chromatin but at the same time the robustness of the overall effects. All tested INO80 mutants generated steady-state patterns (Figure S5B) and differed from WT INO80 in forming aligned arrays in the following ways.

**Figure 5.**
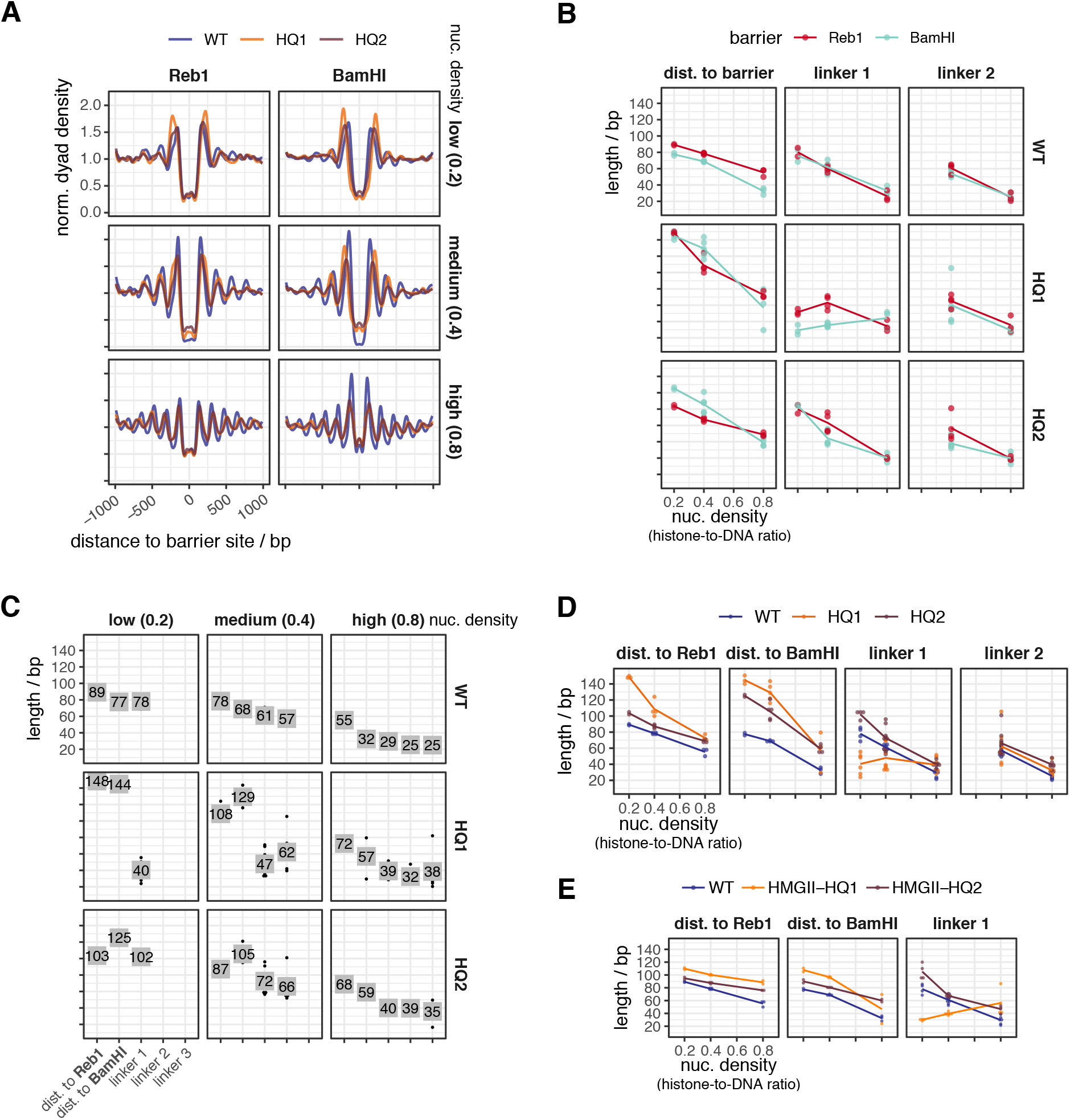
Mutations in the INO80 Arp8 module affect the generation of array features. **(A)** Composite plots as in Figure 2A but for the indicated WT and mutant INO80 complexes and nucleosome densities. **(B) – (D)** As Figure 2C-E, but comparing indicated WT and mutant INO80 complexes. Individual replicates in Figures S5A,B. **(E)** As Figure 2E, but for the indicated WT and mutant INO80 complexes.

**Figure 6.**
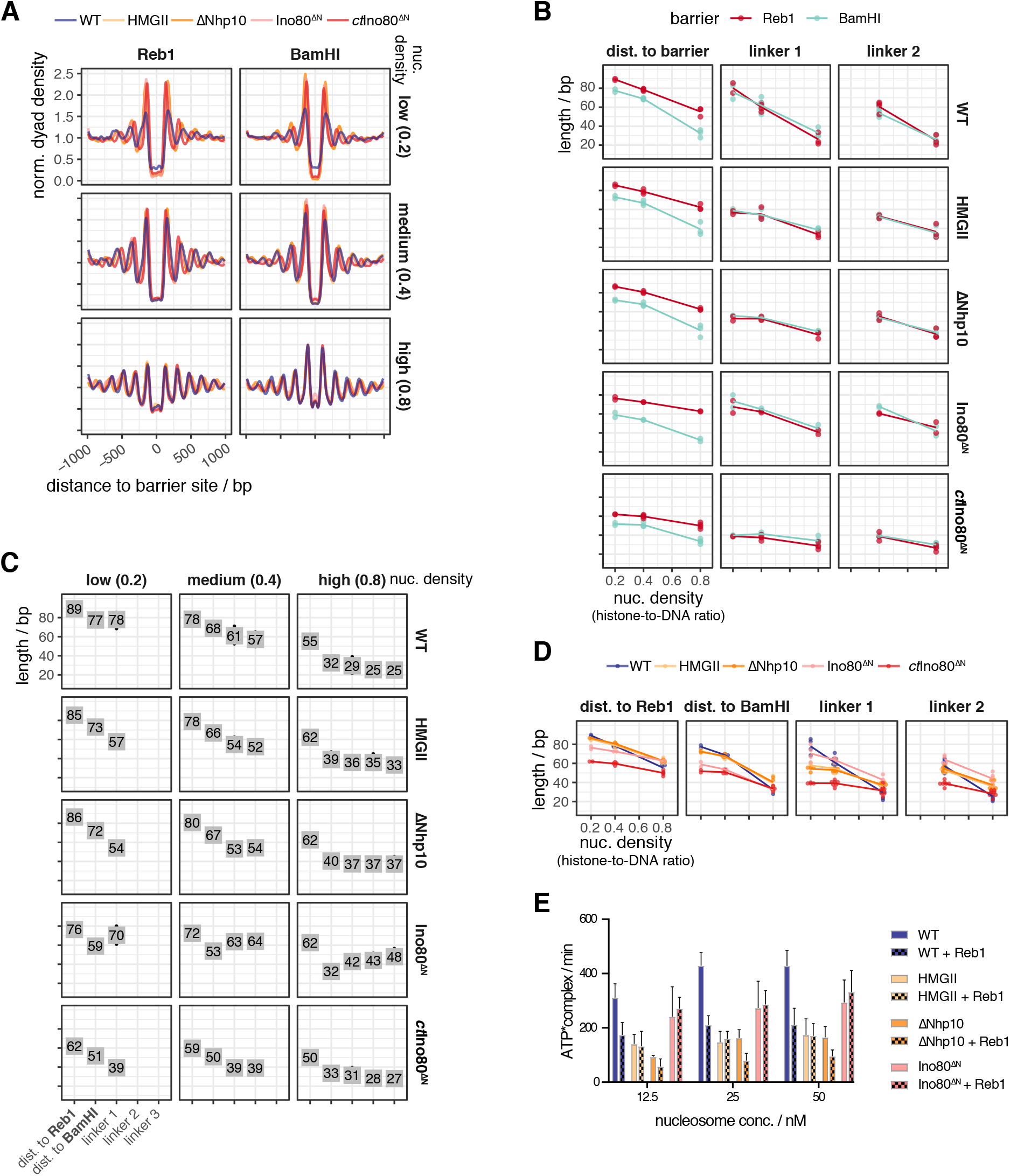
Mutations in the INO80 Nhp10 module affect the generation of array features. **(A) – (D)** As Figure 5A-D but for the indicated WT and mutant INO80 complexes. Individual replicates in Figure S5A. **(E)** Mononucleosome-stimulated ATPase activities for the indicated nucleosome concentrations and INO80 complexes, and respective equimolar Reb1 concentrations. The extranucleosomal DNA of the 601 mononucleosome contained a Reb1 site at 70 bp distance from the 601 sequence and Reb1 was included as indicated.

First, all mutants, besides the HQ1/2 mutant, which was almost inactive (Figure S5A), as expected (Knoll et al., 2018), generated phased regular arrays, but with varying effectiveness and altered distance to one or both barrier types and/or linker lengths compared to WT INO80 (Figures 5D,E, 6D). This revealed that also INO80 has a ruler, to which both the Arp8 and the Nhp10 module contribute.

Second, the HQ1 showed stronger effects than the HQ2 mutation (Figure 5D). Both increased the distances to both barriers. While HQ2 increased linker length at all densities, HQ1 gained clamping activity, i.e., linker length hardly depended on nuclesome density. Both mutations uncoupled distance to DNA ends from linker lengths, in contrast to WT INO80 (Figure 2D,E). Only for HQ1, linker 1 length depended on barrier type (Figure 5B). We concluded that the Arp8 module, especially via HSAα1 helix-DNA interactions, is threefold involved in spacing, alignment to barrier and responding to nucleosome density.

Third, the Nhp10 module subunits contributed to the ruler mainly through the HMG box of Nhp10 as the respective point mutations (HMGII mutant) mimicked the effects upon lack of all Nhp10 module subunits (ΔNhp10 mutant) (Figure 6C,D). With these mutations, distances to both barriers were not much affected, but linker length depended less on density, i.e., clamping was gained, similar to the HQ1 mutation. Effects of the combined HMGII-HQ1 and –HQ2 mutations were dominated by the HQ mutations, but with reduced effects on distance to barriers (Figure 5E). Even though the Nhp10 HMG box was a prime candidate for sensing extranucleosomal DNA, its contribution was minor compared to the HSA helix contribution.

Fourth, the INO80^ΔN^ mutation affected the distance to Reb1 and even more to DNA ends, but gained clamping less strongly than the HMGII or ΔNhp10 mutations (Figure 6D). The INO80^ΔN^ mutant lacked the complete Nhp10 module, but also the Ino80 ATPase N-terminus and Taf14 (Figure S1A), which may account for the differential effects.

Fifth, the INO80^ΔN^, HQ1 and HQ2 mutations most drastically affected distance to BamHI sites, but in opposite ways (Figures 5C,D, 6C,D).

### Effects on nucleosome stimulated ATPase activity versus on ruler function are not strictly coupled

The WT INO80 ATPase activity is stimulated by nucleosomes and inhibited about twofold in the presence of Reb1 (accompanying paper Oberbeckmann & Krietenstein et al.). This relative inhibition by Reb1 was not seen or less pronounced for the mutated INO80 complexes (Figure 6E). The ATPase activity of HMGII and ΔNhp10 mutants was similar to that of WT INO80 in the presence of Reb1. The INO80^ΔN^ mutant had intermediate activity. We concluded that all tested mutants were affected both with regard to ATPase activity and with regard to their ruler but that both effects were not strongly coupled.

### *Chaetomium thermophilum* INO80 core complex suggests species-specific ruler

The INO80 core complex of *C. thermophilum*, which we previously used for cryoEM studies (Eustermann et al., 2018), corresponds to the *S. cerevisiae* INO80^ΔN^ mutant as it also lacks its Ino80 ATPase N-terminus. It showed stronger clamping and generated shorter linkers and distances to Reb1 than INO80^ΔN^ at all densities, and much shorter linkers and distances to both barriers than *S. cerevisiae* WT INO80 at low and medium densities (Figure 6B-D). This suggests that INO80’s ruler may be species-specific.

## Discussion

Our study answers one of the oldest questions in chromatin research: what determines the spacing and phasing distances of nucleosome arrays in absolute terms? The solution to this question are ATP dependent remodelers from the ISWI, CHD and INO80 families with spacing activity. These do not only equalize linker lengths but, as we reveal here, bear rulers for setting distances between two adjacent nucleosomes and between nucleosomes and other alignment points.

### Remodeler rulers explain previous *in vivo* observations

Rulers combined with barriers mechanistically explain *in vivo* observations that involved ISW1a, ISW2, Chd1 and INO80 in +1 nucleosome positioning and/or array regularity and phasing (Gkikopoulos et al., 2011; Hennig et al., 2012; Kubik et al., 2019; Ocampo et al., 2016; Parnell et al., 2015; Pointner et al., 2012; van Bakel et al., 2013; Whitehouse et al., 2007; Yen et al., 2012).

The average *S. cerevisiae* linker length of 18 bp (Thomas and Furber, 1976) results from combined contributions of ISW1a and Chd1 (Ocampo et al., 2016). As we show that ISW1a and Chd1 rulers generate linkers of about 20 and 12 bp, respectively, the 18 bp average linker speaks for ISW1a contributing globally more than Chd1. Indeed, lack of Isw1 *in vivo* globally shortened linkers, while lack of Chd1 affected global spacing only mildly (Kubik et al., 2019; Ocampo et al., 2016). Locally, high transcription rate correlates with shorter spacing (Chereji et al., 2018; Ocampo et al., 2016), which points to increased Chd1 contribution, probably due to increased Chd1 recruitment by elongating RNA polymerase (Simic et al., 2003).

Remodeler-specific rulers explain how ISW1a, ISW2 and INO80 affect +1 nucleosome positioning *in vivo* (Kubik et al., 2019; Parnell et al., 2015; Whitehouse et al., 2007; Yen et al., 2012) and *in vitro* (Krietenstein et al., 2016), especially in combination with RSC. RSC and SWI/SNF are the only yeast remodelers that disassemble nucleosomes (Clapier and Cairns, 2009; Clapier et al., 2017), particularly at promoter NDRs (Badis et al., 2008; Brahma and Henikoff, 2019; Ganguli et al., 2014; Hartley and Madhani, 2009; Kubik et al., 2019; Kubik et al., 2018; Parnell et al., 2008; Rawal et al., 2018; van Bakel et al., 2013; Wippo et al., 2011). By definition, a promoter NDR has low nucleosome density. Therefore, remodeler rulers will set distances to NDR-bound barriers as measured here at low or medium nucleosome density. *In vivo* distances between Reb1 and +1 nucleosomes are 60-80 bp (Figure S3B, (Rhee and Pugh, 2011)), which are within remodeler-specific distances to Reb1 at medium or low density (81-86 bp for INO80, 70-74 bp for ISW2, 58-60 bp for ISW1a). ISW2 and INO80 contribute more to +1 nucleosome positioning *in vivo* than ISW1a (Kubik et al., 2019) as their long rulers are more suited for setting long distances across NDRs. Conversely, the short Chd1-ruler hardly contributes to +1 positioning *in vivo* (Kubik et al., 2019; Ocampo et al., 2016; van Bakel et al., 2013). These different ruler characteristics explain why ISW1a and Chd1 are mainly involved in spacing nucleosomes into densely packed arrays and why ISW2 and INO80 mainly use their ruler for +1 alignment at NDRs *in vivo*. This resolves the conundrum (Krietenstein et al., 2016) why yeast has two remodelers, INO80 and ISW2, that seemingly generate “too wide” spacing compared to average *in vivo* spacing. We do not preclude that other mechanisms, like recruitment via histone modifications or transcription factors, also affect where each remodeler is active.

### Functional and structural identification of remodeler rulers

The protein ruler model was first proposed for ISW1a (Yamada et al., 2011). It suggested that ISW1a shortens the linker until its ruler contacts the neighboring nucleosome, but did not conceptualize why this would lead to a stable nucleosome position. We built on and expanded this model, identified remodeler rulers via their functionality and pinpointed the INO80 ruler also in structural terms. On the functional level, a ruler is revealed if

a. the same remodeler generates the same phasing and/or spacing distances although it works on chromatin with varying nucleosome density (clamping activity), or
b. different remodelers/different mutant versions of the same remodeler generate different phasing and/or spacing distances although they all work on the same chromatin (remodeler-specific phasing/spacing).

For the INO80 complex, we found that the Nhp10 module, especially the Ino80 N-terminus, as well as the Arp8 module, especially the Ino80-HSA-helix, contributed to the ruler function. Lack of the Ino80 N-terminus, concomitant with lacking the Nhp10 module, allowed INO80, e.g., to slide nucleosomes closer to DNA ends, maybe for steric reasons, while impaired DNA traction during remodeling due to compromised Ino80-HSA helix-DNA interactions had the opposite effect. It remains to be elucidated how exactly such modules within the multi-subunit organisation relay barrier information to the core ATPase.

### Remodeler rulers regulate nucleosome sliding direction bias in response to nucleosome environment

We propose an overarching framework for this relay that amounts to a widely applicable remodeler ruler principle (Figure 7). A remodeler may slide a nucleosome either to the left or to the right from a given position. If there is no bias for sliding in either direction, the nucleosome will experience a random walk along the DNA (regions C in in three hypothetical examples Figure 7A). Net nucleosome movement in one direction (Gangaraju and Bartholomew, 2007; Langst et al., 1999; McKnight et al., 2011; Stockdale et al., 2006; Udugama et al., 2011; Yang et al., 2006; Zhou et al., 2018) requires an overall sliding direction bias in this direction. We conceptualize a remodeler ruler as a remodeler-intrinsic feature that generates an overall sliding direction bias in response to the (epi)genetic information in the environment of the nucleosome that the remodeler is remodeling. The bias may originate from differences, e.g., in binding orientation, ATPase activity, sliding rate or processivity and is regulated by interaction of the ruler with a generalized “barrier”. This may be a GRF, a DSB, a neighboring nucleosome, or a DNA sequence element. Histone modifications/ variants may modulate as well. While the microscopic details may differ for different remodelers and information input, the overall regulation of sliding direction bias by the ruler will share three key elements that constitute the ruler mechanism. First, the ruler has a certain reach (regions A + B in Figure 7A), within which it interacts with the barrier. Second, if the position, from where the remodeler slides the nucleosome, is within region B, the interaction between ruler and barrier biases overall sliding direction towards the barrier (red curve is above green curve), e.g., due to binding energy gained upon orienting the remodeler towards vs. away from the barrier. Third, if the nucleosome is in region A, the ruler-barrier interaction disfavors sliding towards relative to sliding away from the barrier (green curve is above red curve), e.g., because the ruler gets sterically in the way. Our study determined the length of region A for different remodeler and barrier types and conditions. Region B and exact curve shapes will have to be determined in future studies. If these three key elements are met, resulting fluxes lead to steady-state nucleosome placement at a defined position relative to the barrier (stippled vertical arrows throughout Figure 7). This position is a self-stabilizing dynamic equilibrium point (intersection of red and green curves) without sliding direction bias here, but with biases *towards* this point from neighboring positions. This model applies to how a remodeler with ruler stably positions a nucleosome next to a GRF as well as to another nucleosome and therefore explains both spacing and phasing.

**Figure 7.**
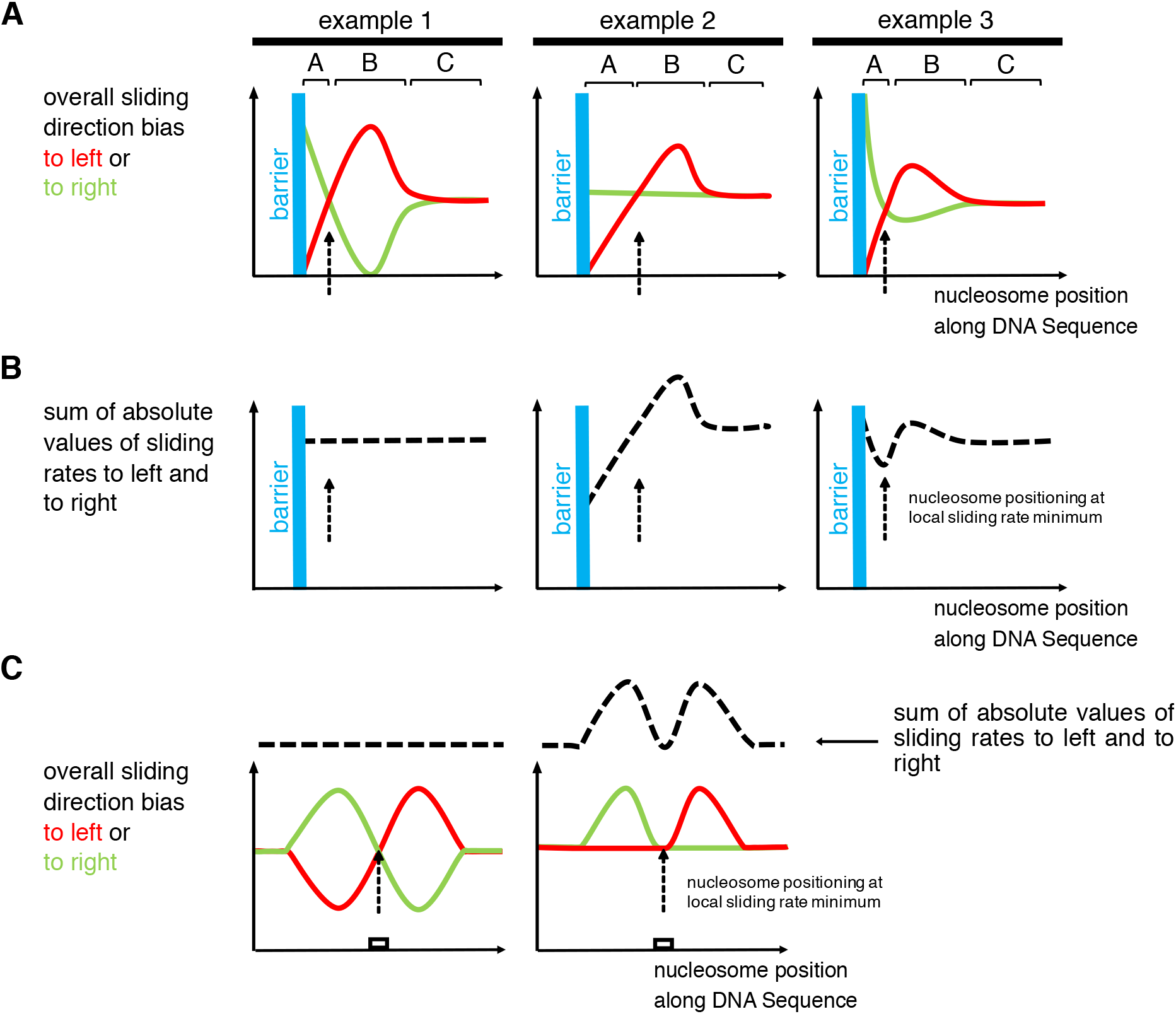
Model for remodeler ruler mechanism. **(A)** Three hypothetical examples for how a remodeler ruler regulates the overall bias of sliding a nucleosome to left (red curves) or to right (green curves) resulting in nucleosome positioning (stippled vertical arrows) in the vicinity of a barrier. **(B)** As panel A but plotting sum of absolute values of sliding rates to the left and to the right (stippled black curves). **(C)** Two hypothetical examples for how a remodeler ruler leads to nucleosome positioning over a DNA site (white box). Symbolics as in panels A and B. For details see text.

It also explains density-independent clamping. As long as a remodeler is processive enough to fortuitously bring nucleosomes into region B of a barrier also at low density, the ruler mechanism will keep the nucleosome at the dynamic equilibrium point. Nonetheless, the model can also accommodate sensing of nucleosome density and barrier type, e.g., if the ruler offers a hierarchy of interaction points that depends on density or barrier type. For example, INO80 may be able to adopt different conformations that may provide different interaction sites and have different footprint sizes, which may explain why INO80 can remodel arrays with just 30 bp linkers despite a measured footprint of >50 bp (Brahma et al., 2018). INO80 mutants showed not concerted but uncoupled effects on distances to Reb1, DNA ends and nucleosomes, even if the same module, like the Nhp10 module, was differentially mutated. Chd1 generated shorter linker lengths (12-16 bp) than distances to DNA ends or Reb1 (35-40 bp). For Chd1, Reb1 may be a “hard” barrier while nucleosomes are “soft” barriers as they are partially “invaded” by the ruler. Indeed, Chd1 partially unwraps nucleosomal DNA (Farnung et al., 2017). The way how different remodeler rulers interact with different barriers requires clarification, and we outline our model (Figure 7) in terms of extension-less point particles, but actual footprints have to be taken into account. The model is fully compatible with the ruler, i.e., the DNA binding domain (DBD) of Chd1 (McKnight et al., 2011) or *Drosophila* ACF (Yang et al., 2006), introducing bias via sensing extranucleosomal DNA length. Indeed, differently long extranucleosomal DNA in mono- or oligonucleosome sliding assays amounts to different distances to barriers like DNA ends or other nucleosomes. Our model is fully consistent with previous data and models but offers an alternative interpretation and is more widely applicable, e.g., to stable nucleosome positioning at only one barrier and not only in-between two barriers.

We introduced our model in terms of overall sliding direction bias. More specifically, the model may refer to differential regulation of sliding rates, i.e., the y-axis in Figure 7A could correspond to “overall sliding rate to the left or to the right”. If sliding rates are reciprocally regulated (example 1, Figure 7A), the sum of absolute sliding rate values is constant at each position (Figure 7B), but not upon asymmetric regulation of sliding direction (examples 2 and 3, Figure 7A). As special case (example 3, Figure 7A,B), the dynamic equilibrium point may correspond to a minimum of absolute sliding rate. This case corresponds to the “kinetic release” model (Manelyte et al., 2014; Rippe et al., 2007), which posits that remodelers position nucleosomes at sites where the nucleosome is the (locally) poorest substrate for remodeling.

### Ruler-regulated sliding: the unifying principle for nucleosome positioning by remodelers

As nucleosome positions are defined by the DNA sequence bound by the histone octamer, all mechanisms that generate consistent nucleosome positions across many genome copies, must select certain DNA sequences in competition with other sequences. As shown here and in the accompanying paper (Oberbeckmann & Krietenstein et al.), remodelers may mediate this selection in two ways. On the one hand, a remodeler may directly choose a sequence, e.g., INO80 turns DNA shape features into +1 nucleosome positions at promoters (accompanying paper Oberbeckmann & Krietenstein et al.) On the other hand, a remodeler ruler may place a nucleosome at a ruler-determined distance to a barrier, e.g., ISW2 aligns nucleosomes to Reb1 and generates a regular array by aligning a second nucleosome to the first and so on. In the former case, the resulting nucleosomal sequence is directly selected for its sequence features, while in the latter case, it is indirectly selected without regards for its sequence features but merely for its position relative to the barrier, as we show here by using Reb1 sites in *S. pombe* and *E. coli* genomes.

Our ruler model unifies these positioning mechanisms. The generalized barrier also encompasses DNA sequence elements, with which a remodeler ruler interacts such that sliding direction bias is regulated (Figure 7C). This explains observations for hybrid Chd1 remodelers where the Chd1 DBD was replaced with heterologous sequence-specific DBDs (Donovan et al., 2019; McKnight et al., 2011; McKnight et al., 2016). Such hybrid Chd1 remodelers slide nucleosomes faster towards the cognate site of the heterologous DBD, if it was in reach of this site, until the nucleosome became positioned on the site. In our model, the heterologous DBD is a remodeler ruler. As a DNA sequence element as barrier is no hindrance for nucleosome sliding, the remodeler may slide the nucleosome onto this site. This prevents ruler binding to the site, abolishes the increase in sliding rate linked to ruler binding and makes a nucleosome on the cognate site a poorer nucleosome sliding substrate than at neighboring positions (Figure 7C, right), which corresponds to the kinetic release model as noted (McKnight et al., 2011). Our model now adds that sliding from neighboring positions will always (within ruler reach) convene at the cognate site and stabilize this position, even if there is no local sliding rate minimum, as long as the ruler regulates sliding direction bias according to the three key elements outlined above (Figure 7C, left). As our INO80 mutations differently affected nucleosome positioning via DNA shape (accompanying paper Oberbeckmann & Krietenstein et al.) vs. relative to Reb1 vs. DNA ends vs. nucleosomes, the ruler elements seem to be multilayered and maybe linked to different structural conformations. For example, the INO80 conformation required for aligning nucleosomes at high density may not be compatible with positioning +1 nucleosomes via DNA shape. Nonetheless, as the Ino80-HSA mutations affected nucleosome positioning both via DNA shape (accompanying paper Oberbeckmann & Krietenstein et al.) and relative to barriers, the Ino80-HSA domain is a functionally crucial part of to the INO80 ruler.

*In vivo* there are many ways that may regulate nucleosome positioning by remodelers, e.g., by recruitment, by architectural factors, by nucleosome density fluctuations or by histone variants and modifications, possibly in the context of elongating polymerases. Nonetheless, we expect that the regulation of nucleosome sliding direction bias via built-in sensing and processing of information in the nucleosome environment, i.e., a remodeler ruler, will be at the heart of each nucleosome positioning mechanism.

## Acknowledgments

We thank Kevin Schall, Nils Krietenstein and Maria Walker for providing purified proteins, Michael Wolff and Ulrich Gerland for helpful discussions of nucleosome positioning mechanisms, Stefan Krebs and Helmut Blum at the Laboratory for Functional Genome Analysis (LAFUGA, Gene Center, LMU München) for high throughput sequencing, and Sigurd Braun for access to and help with the plating robot. This study was funded by the German Research Foundation (SFB1064 to P.K.; SFB860, SPP1935, and EXC 2067/1-390729940 to P.C.), and the European Research Council (grant agreement No 693023 to P.C.).

## Author contributions

Conceptualization: PK, SE, EO, KPH; Data curation: EO, VN; Formal analysis: EO, VN; Funding acquisition, Project administration, Supervision: PK, KPH, SE, PC, CLP; Investigation: EO, VN, SW, LF, MM, AS; Methodology: EO, VN, SE, SW, LF, PK; Validation: EO, VN, MM, AS, SW, LF, PK, SE; Visualization: EO, VN, PK, SW, LF; Writing original draft: PK, SE, EO; Writing – review & editing: PK, SE, EO, VN, KPH, LF, SW, CLP, PC.

## Methods

### Organisms as source for materials used in experiments

The pGP546 yeast genomic plasmid library was expanded from the clonal plates provided by Open Biosystems. For generation of genomic plasmid libraries, the *S. pombe* strain Hu0303 (Ekwall group) and *E. coli* strain (ATCC 11303 strain, 14380, Affymetrix) were used.

INO80 wild-type and mutant complexes, Chd1 and FACT were expressed in *Trichoplusiani insect cells. Spodoptera frugiperda sf21* insect cells were used for virus production. The *Saccharomyces cerevisiae* ISW1a and Fun30 remodelers were purified from the correspondingly TAP-tagged yeast strains, Ioc3-TAP, Fun30-Tap, as provided by Open Biosystems. Yeast ISW2 was purified from strain YTT480 (ISW2-2xFLAG, Tsukiyama et al., 1999). Reb1 was purified from *E. coli* BL21 (DE3) cd+ cells. The *Drosophila* embryo histones were prepared from the *Drosophila melanogaster* strain OregonR.

### Embryonic *D. melanogaster* histones, whole-genome plasmid libraries and salt gradient dialysis

#### Embryonic *D. melanogaster* histone purification

The preparation of embryonic *D. melanogaster* histones octamers was carried out as described in Krietenstein et al. 2012 and Simon and Felsenfeld, 1979. Briefly, 50 g of 0-12 hours old *D. melanogaster* embryos were dechorionated in 3 % sodium hypochlorite, washed with dH_2_0 and resuspended in 40 mL lysis-buffer (15 mM K·HEPES pH 7.5, 10 mM KCl, 5 mM MgCl_2_, 0.1 mM EDTA, 0.5 mM EGTA, 1 mM DTT, 0.2 mM PMSF, 10 % glycerol). Embryos were homogenized (Yamamoto homogenizer), filtered through cloth and centrifuged at 6,500 *g* for 15 min. Nuclei (brownish light pellet) were washed 3 times with 50 mL sucrose-buffer (15 mM K·HEPES pH 7.5, 10 mM KCl, 5 mM MgCl_2_, 0.05 mM EDTA, 0.25 mM EGTA, 1 mM DTT, 0.2 mM PMSF, 1.2 % sucrose) and resuspended in 30 mL sucrose-buffer containing 3 mM CaCl_2_. To obtain mononucleosomes, nuclei were incubated for 10 min at 26 °C with 6250 Units MNase (Sigma-Aldrich). Reaction was stopped with 10 mM EDTA, nuclei were pelleted and resuspended in 6 mL TE (10 mM Tris·HCl pH 7.6, 1 mM EDTA) containing 1 mM DTT and 0.2 mM PMSF followed by 30 to 45 min of rotation at 4 °C. Nuclei were centrifuged for 30 min at 15,300 *g* at 4 °C. Solubilized mononucleosomes are found in the supernatant, which was applied to a pre-equilibrated hydroxyapatite column. After washing the hydroxyapatite column with 0.63 M KCl, histone octamers were eluted with 2 M KCl, concentrated and stored in 50 % glycerol and 1x Complete (Roche) protease inhibitors without EDTA at −20 °C.

#### Whole-genome plasmid library expansion

The *S. cerevisiae* genomic plasmid library (pGP546) was originally described in Jones et al. 2008 and purchased as a clonal glycerol stock collection from Open Biosystems. Library expansion was carried out via a Singer ROTOR plating machine (Singer Instruments) (8-12 rounds, 3 replicas). After 16 hours, colonies were combined into 3×2 L of LB medium containing 50 μg/mL kanamycin and grown for 4 hours. Cells were harvested and subjected to Plasmid Giga Preparation (PC 10 000 Kit, Macherey&Nagel).

For *S. pombe* and *E. coli* plasmid library generation, genomic *S. pombe* (Hu0303) and *E. coli* (type B cells, ATCC 11303 strain, 14380, Affymetrix) DNA was fragmented by a limited SauIIIA or AluI digest. Fragmented DNA was ligated into pJET1.2 vector (ThermoFisher Scientific) and transformed into electrocompetent DH5α cells. Cells were plated on LB plates containing 100 μg/mL ampicillin, grown for 16 – 20 hours, combined in LB medium containing 100 μg/mL ampicillin and grown for another 4 hours. Plasmids was extracted with Plasmid Mega Preparation Kit (PC 2000 Kit, Macherey&Nagel).

#### Salt gradient dialysis (SGD)

For low, medium and high assembly degrees, 10 μg of plasmid library DNA (*S. cerevisiae*, *S. pombe* or *E. coli)* was mixed with ~2, 4 or 8 μg of *Drosophila* embryo histone octamers, respectively, in 100 μl assembly buffer (10 mM Tris·HCl, pH 7.6, 2 M NaCl, 1 mM EDTA, 0.05 % IGEPAL CA630, 0.2 μg BSA). Samples were transferred to Slide-A-lyzer mini dialysis devices, which were placed in a 3 L beaker containing 300 mL of high salt buffer (10 mM Tris·HCl pH 7.6, 2 M NaCl, 1 mM EDTA, 0.05 % IGEPAL CA630, 14.3 mM β-mercaptoethanol), and dialyzed against a total of 3 L low salt buffer (10 mM Tris·HCl pH 7.6, 50 mM NaCl, 1 mM EDTA, 0.05 % IGEPAL CA630, 1.4 mM β-mercaptoethanol) added continuously via a peristaltic pump over a time course of 16 h while stirring. β-mercaptoethanol was added freshly to all buffers. After complete transfer of low salt buffer, samples were dialyzed against 1 L low salt buffer for 1 h at room temperature. DNA concentration of the SGD chromatin preparations was estimated with a DS-11+ spektrophotometer (Denovix) and could be stored at 4 °C for several weeks. To estimate the extent of the assembly degree, an aliquot of the sample was subjected to MNase digestion (as described below) for MNase-ladder read out.

### Purifications of chromatin remodeling enzymes

#### Expression and purification of INO80 complex and respective mutants

Exact strategy for recombinant expression of *S. cerevisiae* INO80 complex in insect cells and complex purification is described in the accompanying paper Krietenstein et al. Briefly, MultiBac technology (Trowitzsch et al., 2010) was applied to generate two baculoviruses carrying coding sequences for *S. cerevisiae* Ino80 (2xFlag), Rvb1, Rvb2, Arp4, Arp5-His, Arp8, Actin, Taf14, Ies1, Ies2, Ies3, Ies4, Ies5, Ies6 and Nhp10 which were subcloned into pFBDM vectors and sequence verified by Sanger Sequencing (GATC Services at Eurofins Genomics). High Five (Hi5) insect cells (BTI-TN-5B1-4 Invitrogen) were co-infected with two or three baculoviruses 1/100 (v/v) each for expression purposes. The recombinantly expressed INO80 complex and respective INO80 mutant complexes were purified from insect cells according to (Tosi, Haas et al. 2013), which resulted in a pure and monodisperse sample. Shortly, cells were resuspended in lysis buffer (50 mM Tris·HCl pH 7.9, 500 mM NaCl, 10 % glycerol, 1 mM DTT, SIGMAFASTTM protease inhibitor cocktail), sonified (Branson Sonifier, 3x 20 s with 40 % duty cycle and output control 3-4) and cleared by centrifugation (Sorvall Evolution RC, SS34 rotor, 15,000 g). The supernatant was incubated with anti-Flag M2 Affinity Gel (Sigma-Aldrich) and centrifuged for 15 min at 1,000 *g* and 4 °C. The anti-Flag resin was washed with buffer A (25 mM K·HEPES pH 8.0, 500 mM KCl, 10 % glycerol, 0.025 mM IGEPAL CA630, 4 mM MgCl_2_, 1 mM DTT) and buffer B (25 mM K·HEPES pH 8.0, 200 mM KCl, 10 % glycerol, 0.02 mM IGEPAL CA630, 4 mM MgCl_2_, 1 mM DTT). Recombinant INO80 complex was eluted with buffer B containing 1.6 mg Flag Peptide (Sigma-Aldrich). Anion exchange chromatography (MonoQ 5/50 GL, GE Healthcare) was used for further purification which resulted in a monodisperse and clear INO80 complex. Using standard cloning techniques, three INO80(2xFlag) HSA domain mutants (HQ1, HQ2, HQ1/2, Figure 4D), one N-terminal deletion mutant (Ino80^ΔN^, deletion of the first 461 amino acids of the N terminus of Ino80) and two INO80 (2xFlag) Nhp10 module mutants (ΔNhp10 (INO80 complex without Ies1, Ies3, Ies5 and Nhp10 but with Ino80 N-terminus) and HMGII (Figure 4G) were generated and integrated into baculoviruses using MultiBac Technology. Expression and purification of mutant INO80 complexes was carried out as described above. The INO80 core complex from *Chaetomium thermophilum* (equivalent to the *S. cerevisiae* N-terminal deletion mutant) was essentially purified as described in Eustermann et al., 2018.

#### Expression and purification of full-length Chd1 and FACT

Hi5 cells (600 mL) were grown in ESF-921 media (Expression Systems) and infected with V1 virus for full-length Chd1 (tagged with a N-terminal 6×His tag, followed by a MBP tag, and a tobacco etch virus protease cleavage site) or FACT (Spt16 carries an N-terminal 6×His tag, followed by an MBP tag, and a tobacco etch virus protease cleavage site) for protein expression. Cells were grown for 72 hours at 72 °C and subsequently harvested by centrifugation (238 *g*, 4 °C, 30 min). Supernatant was discarded and cell pellets resuspended in lysis buffer (300 mM NaCl, 20 mM Na·HEPES pH 7.4, 10 % (v/v) glycerol, 1 mM DTT, 30 mM imidazole pH 8.0, 0.284 μg/mL leupeptin, 1.37 μg/mL pepstatin A, 0.17 mg/mL PMSF, 0.33 mg/mL benzamidine). Resuspended cells were snap frozen and stored at −80 °C.

All protein purifications were performed at 4 °C. Frozen cell pellets were thawed and lysed by sonication. Lysates were cleared using centrifugation (18,000 g, 4 °C, 30 min and 235.000 *g*, 4 °C, 60 min). The supernatant containing Chd1 was filtered with 0.8-μm syringe filters (Millipore) and applied onto a GE HisTrap HP 5 mL (GE Healthcare). The column was washed with 10 column volumes (CV) lysis buffer, 5 CV high salt buffer (1 M NaCl, 20 mM Na·HEPES pH 7.4, 10 % (v/v) glycerol, 1 mM DTT, 30 mM imidazole pH 8.0, 0.284 μg/mL leupeptin, 1.37 μg/mL pepstatin A, 0.17 mg/mL PMSF, 0.33 mg/mL benzamidine), and 5 CV lysis buffer. Chd1 was eluted using a 40-minutes gradient of 0-100 % elution buffer (300 mM NaCl, 20 mM Na·HEPES pH 7.4, 10 % (v/v) glycerol, 1 mM DTT, 500 mM imidazole pH 8.0, 0.284 μg/mL leupeptin, 1.37 μg/mL pepstatin A, 0.17 mg/mL PMSF, 0.33 mg/mL benzamidine). Fractions containing Chd1 were pooled and subjected to dialysis/TEV protease digestion for 16 hours (300 mM NaCl, 20 mM Na·HEPES pH 7.4, 10 % (v/v) glycerol, 1 mM DTT, 30 mM imidazole with 2 mg His6-TEV protease).

The dialyzed sample was again applied to a GE HisTrap HP 5 mL. The flow-through, which contained cleaved tag-less Chd1, was concentrated using an Amicon Millipore 15 mL 50.000 MWCO centrifugal concentrator. The concentrate was applied to a GE S200 16/600 pg size exclusion column in 300 mM NaCl, 20 mM Na·HEPES pH 7.4, 10 % (v/v) glycerol, 1 mM DTT. Fractions containing Chd1 were concentrated to ~100 μM. The sample was aliquoted, snap frozen and stored at −80 °C.

FACT was purified as above with minor modifications. After dialysis, the sample was subjected to a tandem GE HisTrap HP 5mL and GE HiTrap Q 5mL columns combination. After sample application, the columns were washed with lysis buffer and the HisTrap removed. FACT was eluted by applying a high salt buffer gradient from 0-100 % high salt buffer (1 M NaCl, 20 mM Na·HEPES pH 7.4, 10 % (v/v) glycerol, 1 mM DTT, 30 mM imidazole pH 8.0). Fractions with FACT were applied to a GE S200 16/600 pg size exclusion column. Peak fractions with FACT were concentrated to a concentration of ~60 μM, aliquoted, snap frozen, and stored at −80 °C.

#### Expression and purifications of ISW1a, ISW2 and Fun30

Tandem affinity purification of ISW 1a (TAP-Ioc3) and Fun30 (TAP-Fun30) was performed as follows: Cultures were grown in YPD media, harvested cells were washed once with water. The cells were lysed in buffer E (20 mM Na·HEPES pH 7.5, 350 mM NaCl, 10 % glycerol, 0.1 % Tween, and 0.5 mM DTT) and protease inhibitors by grinding in the presence of liquid nitrogen. Lysates were clarified at 40,000 *g* at 4 °C for 1 h. Cleared lysates were incubated with IgG-Sepharose (GE Healthcare) at 4 °C for 2 h and eluted by TEV protease (Invitrogen) cleavage at 4 °C overnight. The elutions were incubated with calmodulin affinity resin (Agilent Technology) in buffer E plus 2 mM CaCl_2_ at 4 °C for 2 h and eluted in buffer E plus 10 mM EGTA.

ISW2 (FLAG-Isw2) was purified as follows: Cleared lysate was incubated with Anti-FLAG M2 affinity gel (Sigma-Aldrich) at 4 °C for 1 h and eluted with 0.1 mg/mL 3X FLAG peptide (Sigma-Aldrich). E-buffer (20 mM Na·HEPES pH 7.5, 350 mM NaCl, 10 % glycerol, 0.1 % Tween, and 0.5 mM DTT) was used during the entire purification.

Purified proteins were concentrated with VIVASPIN concentrators (Sartorius) and dialyzed against E-Buffer with 1 mM DTT. Subunit compositions were confirmed by SDS-PAGE (Figure S1A) and mass spectrometry.

#### Expression and purification of *S. cerevisiae* Reb1

Purification of *S. cerevisiae* Reb 1 was essentially carried out as described in (Krietenstein et al., 2016). Briefly, using BY4741 genomic *S. cerevisiae* DNA the coding sequence for Reb1 was amplified by PCR and cloned into pET21b (Novagen) via InFusion cloning (Clontech) with a Streptavidin tag at the C terminus. Correct sequences were verified via Sanger sequencing (GATC Services at Eurofins Genomics). Expression plasmids were transformed into BL21 (DE3) cd+ cells. Three liters of LB medium supplemented with 600 mg/L ampicillin were inoculated with 200 mL pre-culture. Cells were grown at 37 °C to an OD_600_ of 0.6 (WPA CO8000 cell density meter). Induction was carried out by addition of IPTG to a final concentration of 1 mM. Cells were grown overnight at 18 °C, harvested by centrifugation (3,500 rpm, Sorvall Evolution RC) and stored at −80 °C. Cells were resuspended in lysis buffer (50 mM Tris·HCl pH 7.9, 500 mM NaCl, 7 % glycerol, 1 mM DTT, 7 % sucrose and protease inhibitor (SIGMAFAST^™^ protease inhibitor cocktail, 1:100), sonicated (Branson Sonifier 250, 5 min at 40-50 % duty cycle and output control 4) and cleared by centrifugation (Sorvall Evolution RC, SS34 rotor, 15,000 g). The supernatant was dialyzed over night against 2 L low salt buffer (25 mM K·HEPES pH 8.0, 150 mM KCl, 7 % glycerol, 4 mM MgCl_2_, 1 mM DTT). Cation ion exchange chromatography (HiTrap SP HP 5 mL, elution buffer: 25 mM K·HEPES pH 8.0, 1 M KCl, 7 % glycerol, 4 mM MgCl_2_, 1 mM DTT) followed by size exclusion chromatography (Superdex 200 10/300, buffer: 25 mM K·HEPES pH 8.0, 200 mM KCl, 7 % glycerol, 4 mM MgCl_2_, 1 mM DTT) were used for purification. Peak fractions were analyzed by Coomassie SDS-PAGE. Fractions containing Reb1 were pooled, concentrated and stored at −80 °C.

#### Preparation of mononucleosomes with recombinant human octamers

Canonical human histones were provided by The Histone Source – Protein Expression and Purification (PEP) Facility at Colorado State University. Lyophilized individual human histones were resuspended in 7 M guanidinium chloride, mixed at a 1.2-fold molar excess of H2A/H2B and dialyzed against 2 M NaCl for 16 h. Histone octamers were purified by size exclusion chromatography (HILoad 16/600 Superdex 200 column, GE Healthcare) and stored at −20 °C in 50 % glycerol.

We used fluorescein-labeled Widom 601 DNA (Lowary and Widom 1998) with 80 bp extranucleosomal DNA (80N0 orientation) harboring an in vivo ChIP-Exo verified Reb1 binding site (Rhee and Pugh 2012) of *S. cerevisiae* gene yGL167c (Reb1 binding motif: TTACCC) 64 or 84 bp distant to the 601 sequence. The DNA template (yGL267c_601 and yGL167c_20bp_601) was amplified via PCR, purified by anion exchange chromatography (HiTrap DEAE FF, GE Healthcare) and vacuum concentrated. DNA and assembled histone octamer were mixed in 1.1-fold molar excess of DNA at 2 M NaCl. Over a time-period of 17 h at 4 °C the NaCl concentration was reduced to a final concentration of 50 mM NaCl. Again, anion exchange chromatography was used to purify reconstituted nucleosome core particle (NCP) which were then dialyzed to 50 mM NaCl. NCPs were concentrated to 1 mg/mL and stored at 4 °C.

#### ATPase Assay

As described previously (Eustermann et al., 2018; Knoll et al., 2018), we applied an NADH-based ATPase assay (Kiianitsa et al., 2003) to determine INO80’s ATPase rate. 15 nM INO80 were incubated at 30 °C in a final volume of 50 μl assay buffer (25 mM K·HEPES pH 8.0, 50 mM KCl, 5 mM MgCl_2_, 0.1 mg/mL BSA) with 0.5 mM phosphoenolpyruvate, 2 mM ATP, 0.2 mM NADH and 25 units/mL lactate dehydrogenase/pyruvate kinase (Sigma-Aldrich) to monitor the NADH dependent fluorescence signal in non-binding, black, 384-well plates (Greiner) at an excitation wavelength of 340 nm and an emission wavelength of 460 nm over a 40-min period. We used the Tecan Infinite M1000 (Tecan) plate reader for read out. For all samples, ATPase activity was determined at maximum INO80 WT ATPase activity. ATPase activity was stimulated with 50 nM, 25 nM and 12.5 nM Reb1 site-80N0 mononucleosomes with or without WT Reb1 at indicated concentrations. Using maximal initial linear rates corrected for the buffer blank, we calculated final ATP turnover rates.

#### Genome-wide remodeling reaction

All remodeling reactions, except Chd1-containing reactions, were performed at 30 °C in 100 μL with final buffer conditions of 26.6 mM Na·HEPES pH 7.5, 1 mM Tris·HCl pH 7.6, 85.5 mM NaCl, 8 mM KCl, 10 mM ammonium sulfate, 10 mM creatine phosphate (Sigma-Aldrich), 3 mM MgCl_2_, 2.5 mM ATP, 0.1 mM EDTA, 0.6 mM EGTA, 1 mM DTT, 14 % glycerol, 20 ng/μl creatine kinase (Roche Applied Science). Chd1-containing reactions were performed in 26.6 mM Na·HEPES pH 7.5, 1 mM Tris·HCl pH 7.6, 50 mM NaCl, 10 mM creatine phosphate (Sigma-Aldrich), 3 mM MgCl_2_, 2.5 mM ATP, 0.1 mM EDTA, 0.6 mM EGTA, 1 mM DTT, 14 % glycerol, 20 ng/μl creatine kinase. If called for, 10 nM of remodeling enzyme (but 50 nM Chd1/FACT), 40 nM Reb1 and 20 Units of BamHI (NEB) was added. Before full-length Chd1 (in high-salt buffer) was added to the reaction, it was diluted together with FACT into low salt buffer. For that, full-length Chd1 and purified FACT was mixed in a 1.2:1 molar ratio in high salt buffer (300 mM NaCl, 20 mM Na·HEPES pH 7.4, 10 % (v/v) glycerol, 1 mM DTT), incubated on ice for 5 min and then diluted to 30 mM NaCl final concentration. Remodeling reactions were started by adding 10 μl SGD chromatin corresponding to about 1 μg DNA assembled into nucleosomes and terminated by adding 0.8 Units apyrase (NEB) followed by incubation at 30 °C for 30 min.

#### MNase-seq

After apyrase addition, remodeling reactions were supplemented with CaCl_2_ to a final concentration of 1.5 mM and digested with 100 Units MNase to generate mostly monoucleosomal DNA. Chd1-reaction were incubated with 20 Units MNase to get the same extent of mononucleosomal DNA. 10 mM EDTA and 0.5 % SDS (final concentrations) were added to stop the MNase digest. After proteinase K treatment for 30 min at 37 °C, samples were ethanol precipitated and electrophoresed for 1.5 – 2 h at 100 V using a 1.5 % agarose gel in 1x Tris-acetate-EDTA (TAE) buffer. Mononucleosome bands were excised and purified with PureLink Quick Gel Extraction Kit (ThermoFisher Scientific).

For library preparation, 10-50 ng of mononucleosomal DNA was incubated with 1.25 Units Taq polymerase (NEB), 3.75 Units T4 DNA polymerase (NEB) and 12.5 Units T4-PNK (NEB) in 1x ligation buffer (B0202S, NEB) for 15 min at 12 °C, 15 min at 37 °C and 20 min at 72 °C. To ligate NEBNext Adaptors (0.75 μM final concentration, NEBNext Multiplex Oligos Kit) to the DNA, samples were incubated with T4 DNA ligase (NEB) at 25 °C for 15 min, followed by incubation with 2 Units USER enzyme (NEB) for 10 min at 37 °C. Fragments were purified using 2 volumes AMPure XP beads (Beckman Coulter) and amplified for 8-10 cycles using NEBNext Multiplex Oligos, Phusion High-Fidelity DNA Polymerase (1 U, NEB), deoxynucleotide solution mix (dNTP, 2.5 mM, NEB) and Phusion HF Buffer (1x, NEB). The following protocol was applied for amplification: 98 °C for 30 s, 98 °C for 10 s, 65 °C for 30 s, 72 °C for 30 s with a final amplification step at 72 °C for 5 min. DNA content was assessed by using Qubit dsDNA HS Assay Kit (Invitrogen). PCR reactions were applied to an 1.5 % agarose gel, needed fragment length (~270 bp) was excised and purified via PureLink Quick Gel Extraction Kit (ThermoFisher Scientific). DNA was measured again with Qubit dsDNA HS Assay Kit and diluted to a final concentration of 10 nM (calculation based on the assumption that the DNA fragment length is 272 bp, i. e., 147 bp nucleosomal DNA and 122 bp sequencing adaptor). Diluted samples were pooled according to sequencing reads (~6 Mio reads/ sample). The final pool was quantified with BioAnalyzer (Agilent) and analyzed on an Illumina HiSeq 1500 in 50 bp single-end mode (Laboratory for Functional Genome Analysis, LAFUGA, LMU Munich).

#### Data Processing

Sequencing data was mapped to the SacCer3 (R64), EF2 or *E. coli* strain B (REL606) genome using Bowtie (Langmead et al., 2009). Multiple matches were omitted. After mapping, data was imported into R Studio using GenomicAlignments (Lawrence et al., 2013). Every read was shifted by 73 bp to cover the nucleosome dyad and extended to 50 bp. Genome coverage was calculated and either aligned to in vivo +1 nucleosome positions (Xu et al., 2009), BamHI cut sites, Reb1 SLIM-ChIP hits (Gutin et al., 2018) or Reb1 PWM hits (Badis et al., 2008). Signal was normalized per gene in a 2001 bp window centered on the alignment point. Heatmaps were sorted either by NFR length (distance between in vivo +1 and −1 nucleosome annotated by calling nucleosomes of *in vivo* MNase-seq by Tirosh) or by Reb1 binding score. For the latter, Reb1 SLIM-ChIP data (GSM2916407) was aligned to in vivo +1 nucleosome positions and sorted by signal strength in a 120 bp-window 160 bp upstream of every +1 nucleosome.

For promotor grouping according to Reb1 site orientation, Reb1 SLIM-ChIP hits which contain a PWM site (± 50 bp) and which are located within 400 bp upstream of in vivo +1 nucleosomes were used. Cluster 1 contains promotors where the Reb 1 PWM motif is located on the sense strand and cluster 2, where the Reb1 PWM motif is located on the antisense strand. Cluster 3 contains Reb1 sites at bidirectional promotors.

#### Data Resources

All raw and processed sequencing data generated in this study have been submitted to the NCBI Gene Expression Omnibus (GEO; https://www.ncbi.nlm.nih.gov/geo/) under accession number GSE140614.

## Supplementary Figures

**Figure S1.**
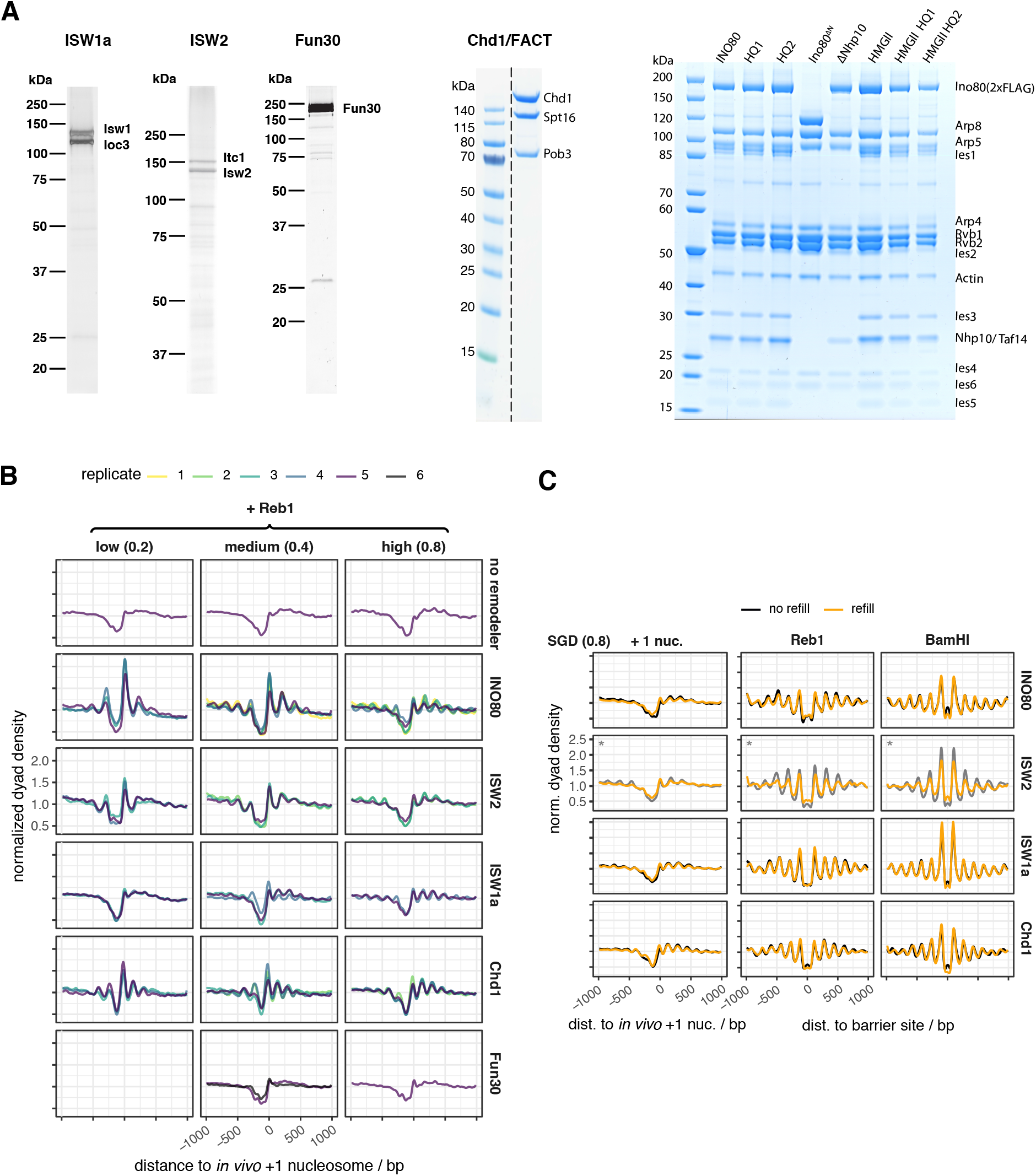
associated with Figures 1 and 2. **(A)** SDS-PAGE analyses of purified remodeler complexes. **(B)** Composite plots as in Figure 1D for individual replicates and the indicated combinations of remodeler, Reb1 and nucleosome density. “no remodeler” denotes absence of remodeler. **(C)** Composite plots aligned at *in vivo* +1 nucleosome positions (left), Reb1 (middle) or BamHI (right) sites for MNase-seq analysis of SGD chromatin assembled at high nucleosome density and incubated with the indicated remodelers as in Figure 2A (no refill) or with doubling remodeler concentration for the second half of incubation time (refill).

**Figure S2.**
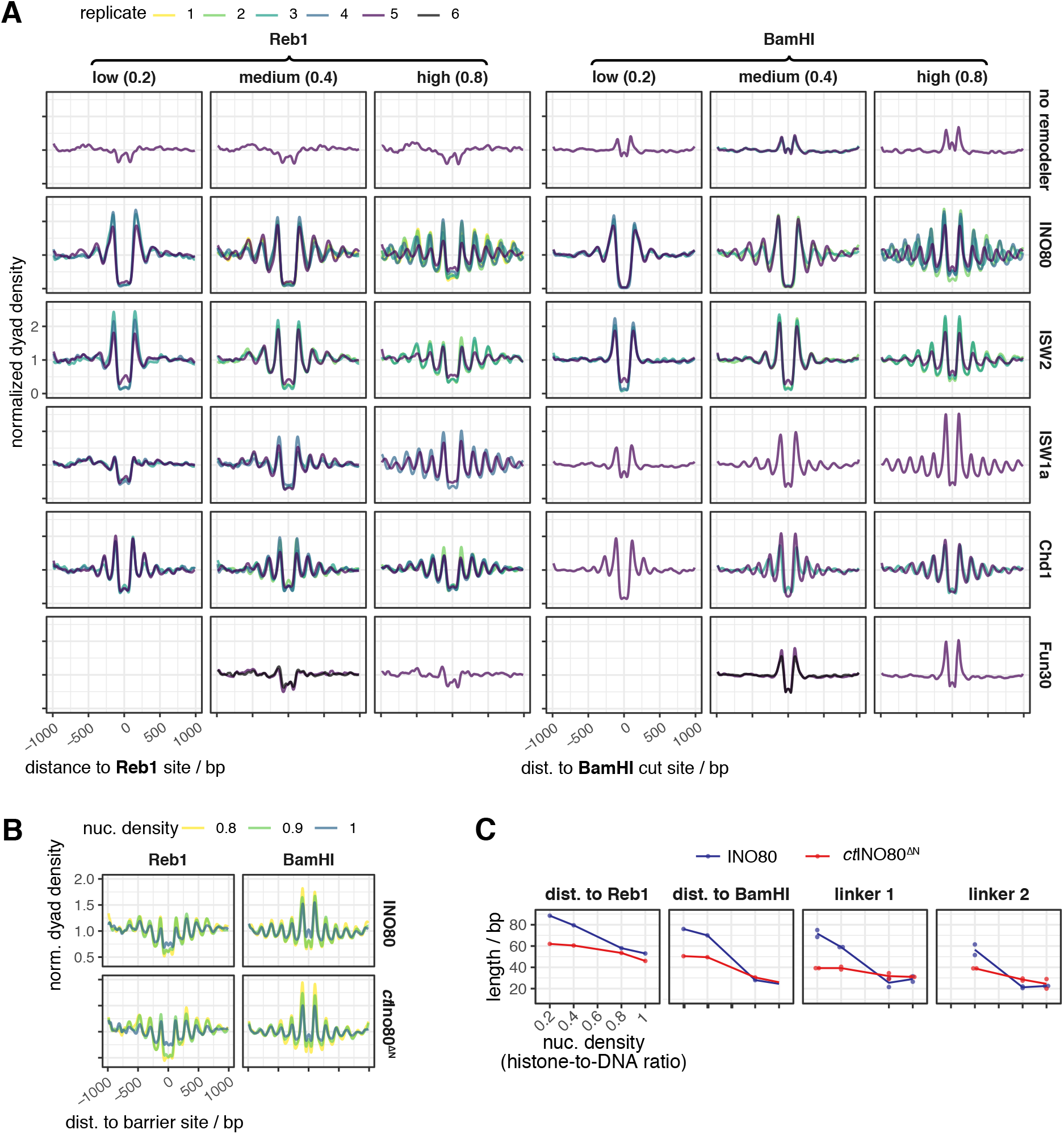
associated with Figure 2. **(A)** As Figure 2A but for individual replicates and indicated combinations of barrier, remodeler and nucleosome density. “no remodeler” denotes absence of remodeler. **(B)** As panel A, but for the indicated nucleosome densities and *S. cerevisiae* WT IN080 complex versus *C. thermophilum* core IN080 complex (ctIN080^ΔN^). **(C)** As Figure 2E, but for the indicated remodelers (as in panel B) and nucleosome densities.

**Figure S3.**
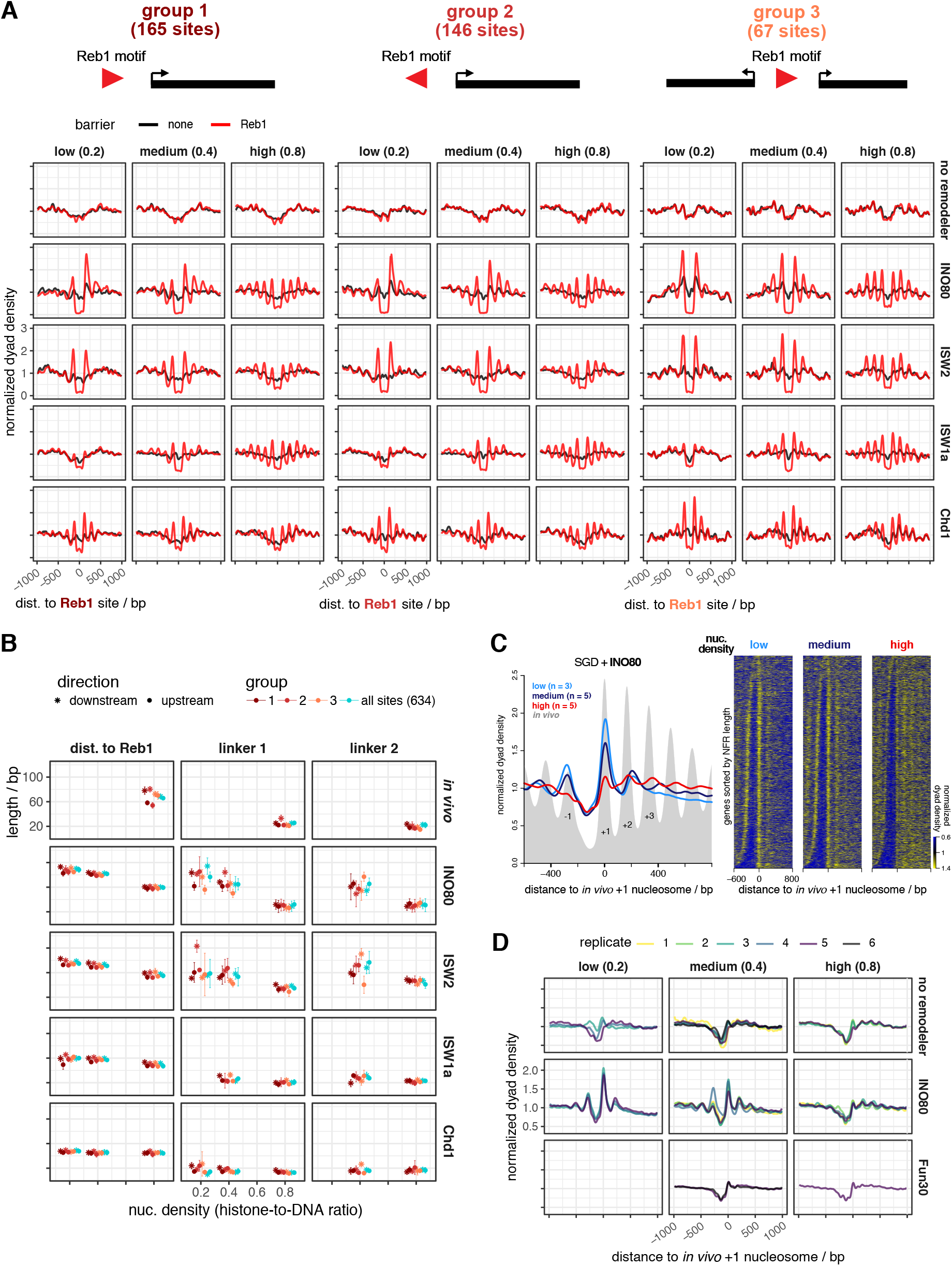
associated with Figure 2. **(A)** As Figure 2A but for indicated combinations of remodeler and nucleosome density. “no remodeler” denotes absence of remodeler. “no remodeler” denotes absence of Reb1. Tracks show merged data from replicates shown in Figures S1B, S2A. Reb1 sites were sorted into groups 1 to 3 (number of sites per group indicated) according to orientation of Reb1 sites relative to one or two genes. **(B)** As Figure 2C, but averages and standard deviation of values for up- and downstream lengths are shown for same groups as in panel A and for all anti-Reb1 SLIM-ChIP sites that contain a Reb1 PWM (light blue). **(C)** Composite plots (left) and heat maps (right) of MNase-seq analysis of *in vivo* chromatin or SGD chromatin reconstituted with the indicated nucleosome density and incubated with recombinant WT INO80 complex. Data are aligned at *in vivo* +1 nucleosome positions and heat maps are sorted from top to bottom by increasing NFR length. Traces with indicated replicate number (n) represent merged data for all replicates. Positions of −1, +1, +2, +3 nucleosomes of the *in vivo* pattern are labeled. **(D)** Composite plots as in panel C, right, but for individual replicates and the indicated remodelers. “no remodeler” denotes absence of remodeler.

**Figure S4.**
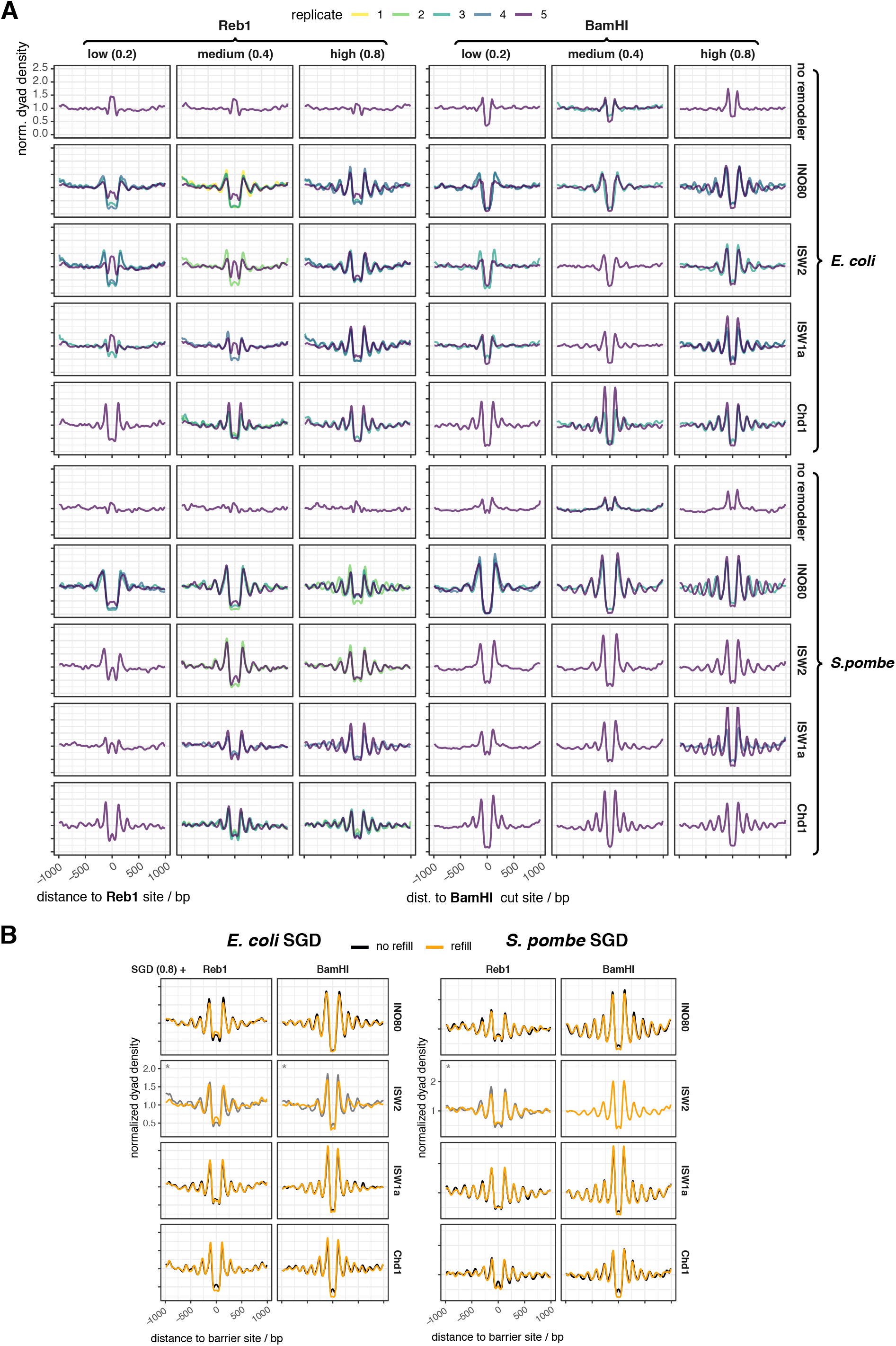
associated with Figure 3. **(A)** As Figure S2A, but for the indicated genomes. Replicate 5 corresponds to the reconstitution with mixed genomes. **(B)** As Figure S2B, but for the indicated genomes.

**Figure S5.**
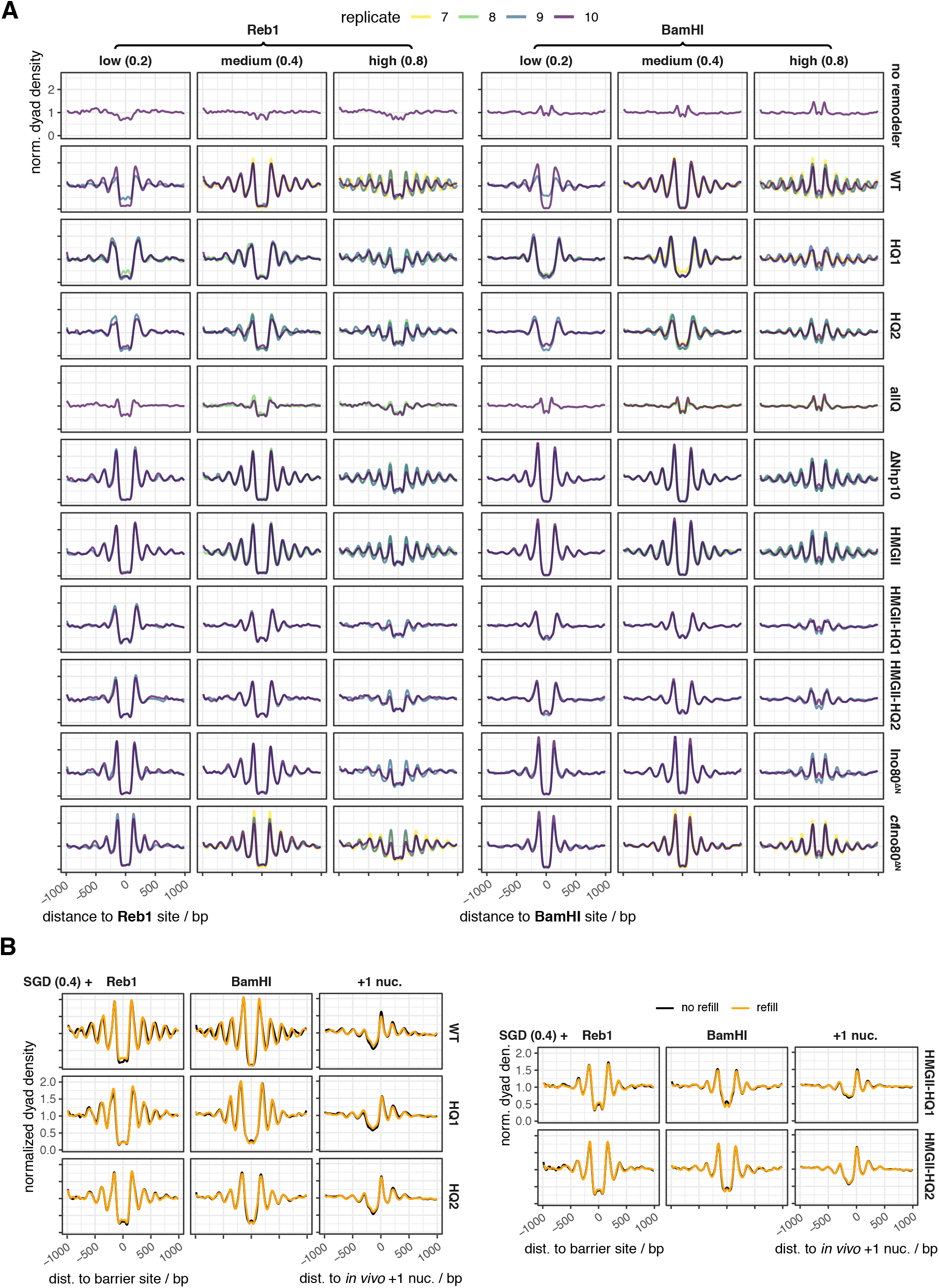
associated with Figures 5, 6. **(A)** As Figure S2A, but for the indicated WT and mutant IN080 remodelers and the *C. thermophilum* IN080 core complex (ctIN080^ΔN^). **(B)** As Figure S2B, but for indicated remodelers.

## References

Awad, S., Ryan, D., Prochasson, P., Owen-Hughes, T., and Hassan, A.H. (2010). The Snf2 homolog Fun30 acts as a homodimeric ATP-dependent chromatin-remodeling enzyme. The Journal of biological chemistry 285, 9477–9484.

Badis, G., Chan, E.T., van Bakel, H., Pena-Castillo, L., Tillo, D., Tsui, K., Carlson, C.D., Gossett, A.J., Hasinoff, M.J., Warren, C.L., et al. (2008). A library of yeast transcription factor motifs reveals a widespread function for Rsc3 in targeting nucleosome exclusion at promoters. Molecular cell 32, 878–887.

Baldi, S., Jain, D.S., Harpprecht, L., Zabel, A., Scheibe, M., Butter, F., Straub, T., and Becker, P.B. (2018a). Genome-wide Rules of Nucleosome Phasing in Drosophila. Molecular cell 72, 661–672.e664.

Baldi, S., Korber, P., and Becker, P.B. (2020). Beads on a string-nucleosome array arrangements and folding of the chromatin fiber. Nature structural & molecular biology 27, 109–118.

Baldi, S., Krebs, S., Blum, H., and Becker, P.B. (2018b). Genome-wide measurement of local nucleosome array regularity and spacing by nanopore sequencing. Nature structural & molecular biology 25, 894–901.

Brahma, S., and Henikoff, S. (2019). RSC-Associated Subnucleosomes Define MNase-Sensitive Promoters in Yeast. Molecular cell 73, 238–249.e233.

Brahma, S., Ngubo, M., Paul, S., Udugama, M., and Bartholomew, B. (2018). The Arp8 and Arp4 module acts as a DNA sensor controlling INO80 chromatin remodeling. Nature communications 9, 3309.

Challal, D., Barucco, M., Kubik, S., Feuerbach, F., Candelli, T., Geoffroy, H., Benaksas, C., Shore, D., and Libri, D. (2018). General Regulatory Factors Control the Fidelity of Transcription by Restricting Non-coding and Ectopic Initiation. Molecular cell 72, 955–969.e957.

Chereji, R.V., Ramachandran, S., Bryson, T.D., and Henikoff, S. (2018). Precise genome-wide mapping of single nucleosomes and linkers in vivo. Genome biology 19, 19.

Clapier, C.R., and Cairns, B.R. (2009). The biology of chromatin remodeling complexes. Annual review of biochemistry 78, 273–304.

Clapier, C.R., Iwasa, J., Cairns, B.R., and Peterson, C.L. (2017). Mechanisms of action and regulation of ATP-dependent chromatin-remodelling complexes. Nature reviews Molecular cell biology 18, 407–422.

Donovan, D.A., Crandall, J.G., Banks, O.G.B., Jensvold, Z.D., Truong, V., Dinwiddie, D., McKnight, L.E., and McKnight, J.N. (2019). Engineered Chromatin Remodeling Proteins for Precise Nucleosome Positioning. Cell reports 29, 2520– 2535.e2524.

Eustermann, S., Schall, K., Kostrewa, D., Lakomek, K., Strauss, M., Moldt, M., and Hopfner, K.P. (2018). Structural basis for ATP-dependent chromatin remodelling by the INO80 complex. Nature 556, 386–390.

Farnung, L., Vos, S.M., Wigge, C., and Cramer, P. (2017). Nucleosome-Chd1 structure and implications for chromatin remodelling. Nature.

Flaus, A., Martin, D.M., Barton, G.J., and Owen-Hughes, T. (2006). Identification of multiple distinct Snf2 subfamilies with conserved structural motifs. Nucleic acids research 34, 2887–2905.

Ganapathi, M., Palumbo, M.J., Ansari, S.A., He, Q., Tsui, K., Nislow, C., and Morse, R.H. (2011). Extensive role of the general regulatory factors, Abf1 and Rap1, in determining genome-wide chromatin structure in budding yeast. Nucleic acids research 39, 2032–2044.

Gangaraju, V.K., and Bartholomew, B. (2007). Dependency of ISW1a chromatin remodeling on extranucleosomal DNA. Molecular and cellular biology 27, 3217–3225.

Ganguli, D., Chereji, R.V., Iben, J.R., Cole, H.A., and Clark, D.J. (2014). RSC-dependent constructive and destructive interference between opposing arrays of phased nucleosomes in yeast. Genome research 24, 1637–1649.

Gkikopoulos, T., Schofield, P., Singh, V., Pinskaya, M., Mellor, J., Smolle, M., Workman, J.L., Barton, G.J., and Owen-Hughes, T. (2011). A role for Snf2-related nucleosome-spacing enzymes in genome-wide nucleosome organization. Science (New York, NY) 333, 1758–1760.

Gossett, A.J., and Lieb, J.D. (2012). In vivo effects of histone H3 depletion on nucleosome occupancy and position in Saccharomyces cerevisiae. PLoS genetics 8, e1002771.

Gutin, J., Sadeh, R., Bodenheimer, N., Joseph-Strauss, D., Klein-Brill, A., Alajem, A., Ram, O., and Friedman, N. (2018). Fine-Resolution Mapping of TF Binding and Chromatin Interactions. Cell reports 22, 2797–2807.

Hartley, P.D., and Madhani, H.D. (2009). Mechanisms that specify promoter nucleosome location and identity. Cell 137, 445–458.

Hennig, B.P., Bendrin, K., Zhou, Y., and Fischer, T. (2012). Chd1 chromatin remodelers maintain nucleosome organization and repress cryptic transcription. EMBO reports 13, 997–1003.

Ito, T., Bulger, M., Pazin, M.J., Kobayashi, R., and Kadonaga, J.T. (1997). ACF, an ISWI-containing and ATP-utilizing chromatin assembly and remodeling factor. Cell 90, 145–155.

Jaiswal, R., Choudhury, M., Zaman, S., Singh, S., Santosh, V., Bastia, D., and Escalante, C.R. (2016). Functional architecture of the Reb1-Ter complex of Schizosaccharomyces pombe. Proceedings of the National Academy of Sciences of the United States of America 113, E2267–2276.

Knoll, K.R., Eustermann, S., Niebauer, V., Oberbeckmann, E., Stoehr, G., Schall, K., Tosi, A., Schwarz, M., Buchfellner, A., Korber, P., et al. (2018). The nuclear actin-containing Arp8 module is a linker DNA sensor driving INO80 chromatin remodeling. Nature structural & molecular biology 25, 823–832.

Kornberg, R.D. (1974). Chromatin structure: a repeating unit of histones and DNA. Science (New York, NY) 184, 868–871.

Kornberg, R.D., and Lorch, Y. (1999). Twenty-five years of the nucleosome, fundamental particle of the eukaryote chromosome. Cell 98, 285–294.

Krietenstein, N., Wal, M., Watanabe, S., Park, B., Peterson, C.L., Pugh, B.F., and Korber, P. (2016). Genomic Nucleosome Organization Reconstituted with Pure Proteins. Cell 167, 709– 721.e712.

Kubik, S., Bruzzone, M.J., Challal, D., Dreos, R., Mattarocci, S., Bucher, P., Libri, D., and Shore, D. (2019). Opposing chromatin remodelers control transcription initiation frequency and start site selection. Nature structural & molecular biology 26, 744–754.

Kubik, S., O’Duibhir, E., de Jonge, W.J., Mattarocci, S., Albert, B., Falcone, J.L., Bruzzone, M.J., Holstege, F.C.P., and Shore, D. (2018). Sequence-Directed Action of RSC Remodeler and General Regulatory Factors Modulates +1 Nucleosome Position to Facilitate Transcription. Molecular cell 71, 89– 102.e105.

Lai, W.K.M., and Pugh, B.F. (2017). Understanding nucleosome dynamics and their links to gene expression and DNA replication. Nature reviews Molecular cell biology 18, 548–562.

Langst, G., Bonte, E.J., Corona, D.F., and Becker, P.B. (1999). Nucleosome movement by CHRAC and ISWI without disruption or trans-displacement of the histone octamer. Cell 97, 843–852.

Lieleg, C., Ketterer, P., Nuebler, J., Ludwigsen, J., Gerland, U., Dietz, H., Mueller-Planitz, F., and Korber, P. (2015). Nucleosome spacing generated by ISWI and CHD1 remodelers is constant regardless of nucleosome density. Molecular and cellular biology 35, 1588–1605.

Luger, K., Mader, A.W., Richmond, R.K., Sargent, D.F., and Richmond, T.J. (1997). Crystal structure of the nucleosome core particle at 2.8 A resolution. Nature 389, 251–260.

Lusser, A., Urwin, D.L., and Kadonaga, J.T. (2005). Distinct activities of CHD1 and ACF in ATP-dependent chromatin assembly. Nature structural & molecular biology 12, 160–166.

McKnight, J.N., Jenkins, K.R., Nodelman, I.M., Escobar, T., and Bowman, G.D. (2011). Extranucleosomal DNA binding directs nucleosome sliding by Chd1. Molecular and cellular biology 31, 4746–4759.

McKnight, J.N., Tsukiyama, T., and Bowman, G.D. (2016). Sequence-targeted nucleosome sliding in vivo by a hybrid Chd1 chromatin remodeler. Genome research 26, 693–704.

Ngo, H.B., Lovely, G.A., Phillips, R., and Chan, D.C. (2014). Distinct structural features of TFAM drive mitochondrial DNA packaging versus transcriptional activation. Nature communications 5, 3077.

Ocampo, J., Chereji, R.V., Eriksson, P.R., and Clark, D.J. (2016). The ISW1 and CHD1 ATP-dependent chromatin remodelers compete to set nucleosome spacing in vivo. Nucleic acids research 44, 4625–4635.

Olins, A.L., and Olins, D.E. (1974). Spheroid chromatin units (v bodies). Science (New York, NY) 183, 330–332.

Olins, D.E., and Olins, A.L. (2003). Chromatin history: our view from the bridge. Nature reviews Molecular cell biology 4, 809–814.

Parnell, T.J., Huff, J.T., and Cairns, B.R. (2008). RSC regulates nucleosome positioning at Pol II genes and density at Pol III genes. The EMBO journal 27, 100–110.

Parnell, T.J., Schlichter, A., Wilson, B.G., and Cairns, B.R. (2015). The chromatin remodelers RSC and ISW1 display functional and chromatin-based promoter antagonism. eLife 4, e06073.

Patel, A., McKnight, J.N., Genzor, P., and Bowman, G.D. (2011). Identification of residues in chromodomain helicase DNA-binding protein 1 (Chd1) required for coupling ATP hydrolysis to nucleosome sliding. The Journal of biological chemistry 286, 43984–43993.

Pointner, J., Persson, J., Prasad, P., Norman-Axelsson, U., Stralfors, A., Khorosjutina, O., Krietenstein, N., Svensson, J.P., Ekwall, K., and Korber, P. (2012). CHD1 remodelers regulate nucleosome spacing in vitro and align nucleosomal arrays over gene coding regions in S. pombe. The EMBO journal 31, 4388–4403.

Rawal, Y., Chereji, R.V., Qiu, H., Ananthakrishnan, S., Govind, C.K., Clark, D.J., and Hinnebusch, A.G. (2018). SWI/SNF and RSC cooperate to reposition and evict promoter nucleosomes at highly expressed genes in yeast. Genes & development 32, 695–710.

Rhee, H.S., and Pugh, B.F. (2011). Comprehensive genome-wide protein-DNA interactions detected at single-nucleotide resolution. Cell 147, 1408–1419.

Rippe, K., Schrader, A., Riede, P., Strohner, R., Lehmann, E., and Langst, G. (2007). DNA sequence-and conformation-directed positioning of nucleosomes by chromatin-remodeling complexes. Proceedings of the National Academy of Sciences of the United States of America 104, 15635–15640.

Satchwell, S.C., Drew, H.R., and Travers, A.A. (1986). Sequence periodicities in chicken nucleosome core DNA. Journal of molecular biology 191, 659–675.

Simic, R., Lindstrom, D.L., Tran, H.G., Roinick, K.L., Costa, P.J., Johnson, A.D., Hartzog, G.A., and Arndt, K.M. (2003). Chromatin remodeling protein Chd1 interacts with transcription elongation factors and localizes to transcribed genes. The EMBO journal 22, 1846–1856.

Smolle, M., Venkatesh, S., Gogol, M.M., Li, H., Zhang, Y., Florens, L., Washburn, M.P., and Workman, J.L. (2012). Chromatin remodelers Isw1 and Chd1 maintain chromatin structure during transcription by preventing histone exchange. Nature structural & molecular biology 19, 884–892.

Stockdale, C., Flaus, A., Ferreira, H., and Owen-Hughes, T. (2006).Analysis of nucleosome repositioning by yeast ISWI and Chd1 chromatin remodeling complexes. The Journal of biological chemistry 281, 16279–16288.

Thomas, J.O., and Furber, V. (1976). Yeast chromatin structure. FEBS letters 66, 274–280.

Torigoe, S.E., Patel, A., Khuong, M.T., Bowman, G.D., and Kadonaga, J.T. (2013). ATP-dependent chromatin assembly is functionally distinct from chromatin remodeling. eLife 2, e00863.

Tsankov, A., Yanagisawa, Y., Rhind, N., Regev, A., and Rando, O.J. (2011). Evolutionary divergence of intrinsic and transregulated nucleosome positioning sequences reveals plastic rules for chromatin organization. Genome research 21, 1851–1862.

Tsukiyama, T., Palmer, J., Landel, C.C., Shiloach, J., and Wu, C. (1999). Characterization of the imitation switch subfamily of ATP-dependent chromatin-remodeling factors in Saccharomyces cerevisiae. Genes & development 13, 686–697.

Udugama, M., Sabri, A., and Bartholomew, B. (2011). The INO80 ATP-dependent chromatin remodeling complex is a nucleosome spacing factor. Molecular and cellular biology 31, 662–673.

Valouev, A., Johnson, S.M., Boyd, S.D., Smith, C.L., Fire, A.Z., and Sidow, A. (2011). Determinants of nucleosome organization in primary human cells. Nature 474, 516–520.

van Bakel, H., Tsui, K., Gebbia, M., Mnaimneh, S., Hughes, T.R., and Nislow, C. (2013). A compendium of nucleosome and transcript profiles reveals determinants of chromatin architecture and transcription. PLoS genetics 9, e1003479.

van Holde, K.E. (1989). Chromatin (New York: Springer).

Varga-Weisz, P.D., Wilm, M., Bonte, E., Dumas, K., Mann, M., and Becker, P.B. (1997). Chromatin-remodelling factor CHRAC contains the ATPases ISWI and topoisomerase II. Nature 388, 598–602.

Whitehouse, I., Rando, O.J., Delrow, J., and Tsukiyama, T. (2007).Chromatin remodelling at promoters suppresses antisense transcription. Nature 450, 1031–1035.

Wiechens, N., Singh, V., Gkikopoulos, T., Schofield, P., Rocha, S., and Owen-Hughes, T. (2016). The Chromatin Remodelling Enzymes SNF2H and SNF2L Position Nucleosomes adjacent to CTCF and Other Transcription Factors. PLoS genetics 12, e1005940.

Wippo, C.J., Israel, L., Watanabe, S., Hochheimer, A., Peterson, C.L., and Korber, P. (2011). The RSC chromatin remodelling enzyme has a unique role in directing the accurate positioning of nucleosomes. The EMBO journal 30, 1277–1288.

Yamada, K., Frouws, T.D., Angst, B., Fitzgerald, D.J., DeLuca, C., Schimmele, K., Sargent, D.F., and Richmond, T.J. (2011). Structure and mechanism of the chromatin remodelling factor ISW 1a. Nature 472, 448–453.

Yan, C., Chen, H., and Bai, L. (2018). Systematic Study of Nucleosome-Displacing Factors in Budding Yeast. Molecular cell 71, 294–305.e294.

Yang, J.G., Madrid, T.S., Sevastopoulos, E., and Narlikar, G.J. (2006). The chromatin-remodeling enzyme ACF is an ATP-dependent DNA length sensor that regulates nucleosome spacing. Nature structural & molecular biology 13, 1078–1083.

Yarragudi, A., Miyake, T., Li, R., and Morse, R.H. (2004). Comparison of ABF1 and RAP1 in chromatin opening and transactivator potentiation in the budding yeast Saccharomyces cerevisiae. Molecular and cellular biology 24, 9152–9164.

Yen, K., Vinayachandran, V., Batta, K., Koerber, R.T., and Pugh, B.F. (2012). Genome-wide nucleosome specificity and directionality of chromatin remodelers. Cell 149, 1461–1473.

Zhang, Y., Moqtaderi, Z., Rattner, B.P., Euskirchen, G., Snyder, M., Kadonaga, J.T., Liu, X.S., and Struhl, K. (2009). Intrinsic histone-DNA interactions are not the major determinant of nucleosome positions in vivo. Nature structural & molecular biology 16, 847–852.

Zhang, Z., Wippo, C.J., Wal, M., Ward, E., Korber, P., and Pugh, B.F. (2011). A packing mechanism for nucleosome organization reconstituted across a eukaryotic genome. Science (New York, NY) 332, 977–980.

Zhou, C.Y., Johnson, S.L., Lee, L.J., Longhurst, A.D., Beckwith, S.L., Johnson, M.J., Morrison, A.J., and Narlikar, G.J. (2018). The Yeast INO80 Complex Operates as a Tunable DNA Length-Sensitive Switch to Regulate Nucleosome Sliding. Molecular cell 69, 677–688.e679.

## References

Badis, G., Chan, E.T., van Bakel, H., Pena-Castillo, L., Tillo, D., Tsui, K., Carlson, C.D., Gossett, A.J., Hasinoff, M.J., Warren, C.L., et al. (2008). A Library of Yeast Transcription Factor Motifs Reveals a Widespread Function for Rsc3 in Targeting Nucleosome Exclusion at Promoters. Molecular Cell 32, 878–887.

Eustermann, S., Schall, K., Kostrewa, D., Lakomek, K., Strauss, M., Moldt, M., and Hopfner, K.-P. (2018). Structural basis for ATP-dependent chromatin remodelling by the INO80 complex. Nature 556, 386–390.

Gutin, J., Sadeh, R., Bodenheimer, N., Joseph-Strauss, D., Klein-Brill, A., Alajem, A., Ram, O., and Friedman, N. (2018). Fine-Resolution Mapping of TF Binding and Chromatin Interactions. Cell Reports 22, 2797–2807.

Jones, G.M., Stalker, J., Humphray, S., West, A., Cox, T., Rogers, J., Dunham, I., and Prelich, G. (2008). A systematic library for comprehensive overexpression screens in Saccharomyces cerevisiae. Nat Meth 5, 239–241.

Kiianitsa, K., Solinger, J.A., and Heyer, W.-D. (2003). NADH-coupled microplate photometric assay for kinetic studies of ATP-hydrolyzing enzymes with low and high specific activities. Anal. Biochem. 321, 266–271.

Knoll, K.R., Eustermann, S., Niebauer, V., Oberbeckmann, E., Stoehr, G., Schall, K., Tosi, A., Schwarz, M., Buchfellner, A., Korber, P., et al. (2018). The nuclear actin-containing Arp8 module is a linker DNA sensor driving INO80 chromatin remodeling. Nat Struct Mol Biol 25, 823–832.

Krietenstein, N., Wippo, C.J., Lieleg, C., and Korber, P. (2012). Genome-Wide In Vitro Reconstitution of Yeast Chromatin with In Vivo-Like Nucleosome Positioning. In Methods in Enzymology, (Elsevier), pp. 205–232.

Krietenstein, N., Wal, M., Watanabe, S., Park, B., Peterson, C.L., Pugh, B.F., and Korber, P. (2016). Genomic Nucleosome Organization Reconstituted with Pure Proteins. Cell 167, 709–721.e12.

Langmead, B., Trapnell, C., Pop, M., and Salzberg, S.L. (2009). Ultrafast and memory-efficient alignment of short DNA sequences to the human genome. Genome Biology 10, R25.

Lawrence, M., Huber, W., Pagès, H., Aboyoun, P., Carlson, M., Gentleman, R., Morgan, M.T., and Carey, V.J. (2013). Software for Computing and Annotating Genomic Ranges. PLOS Computational Biology 9, e1003118.

Simon, R.H., and Felsenfeld, G. (1979). A new procedure for purifying histone pairs H2A + H2B and H3 + H4 from chromatin using hydroxylapatite. Nucleic Acids Res 6, 689–696.

Xu, Z., Wei, W., Gagneur, J., Perocchi, F., Clauder-Münster, S., Camblong, J., Guffanti, E., Stutz, F., Huber, W., and Steinmetz, L.M. (2009). Bidirectional promoters generate pervasive transcription in yeast. Nature 457, 1033–1037.

